# A machine learning and drug repurposing approach to target ferroptosis in colorectal cancer stratified by sex and KRAS

**DOI:** 10.1101/2024.06.24.600340

**Authors:** Hong Yan, Xinyi Shen, Yisha Yao, Sajid A. Khan, Shuangge Ma, Caroline H. Johnson

## Abstract

The landscape of sex differences in Colorectal Cancer (CRC) has not been well characterized with respect to the mechanisms of action for oncogenes such as KRAS. However, our recent study showed that tumors from male patients with KRAS mutations have decreased iron-dependent cell death called ferroptosis. Building on these findings, we further examined ferroptosis in CRC, considering both sex of the patient and KRAS mutations, using public databases and our in-house CRC tumor cohort.

Through subsampling inference and variable importance analysis (VIMP), we identified significant differences in gene expression between KRAS mutant and wild type tumors from male patients. These genes suppress (e.g., *SLC7A11*) or drive (e.g., *SLC1A5*) ferroptosis, and these findings were further validated with Gaussian mixed models. Furthermore, we explored the prognostic value of ferroptosis regulating genes and discovered sex- and KRAS-specific differences at both the transcriptional and metabolic levels by random survival forest with backward elimination algorithm (RSF-BE). Of note, genes and metabolites involved in arginine synthesis and glutathione metabolism were uniquely associated with prognosis in tumors from males with KRAS mutations.

Additionally, drug repurposing is becoming popular due to the high costs, attrition rates, and slow pace of new drug development, offering a way to treat common and rare diseases more efficiently. Furthermore, increasing evidence has shown that ferroptosis inhibition or induction can improve drug sensitivity or overcome chemotherapy drug resistance. Therefore, we investigated the correlation between gene expression, metabolite levels, and drug sensitivity across all CRC primary tumor cell lines using data from the Genomics of Drug Sensitivity in Cancer (GDSC) resource. We observed that ferroptosis suppressor genes such as *DHODH*, *GCH1*, and *AIFM2* in KRAS mutant CRC cell lines were resistant to cisplatin and paclitaxel, underscoring why these drugs are not effective for these patients. The comprehensive map generated here provides valuable biological insights for future investigations, and the findings are supported by rigorous analysis of large-scale publicly available data and our in-house cohort. The study also emphasizes the potential application of VIMP, Gaussian mixed models, and RSF-BE models in the multi-omics research community. In conclusion, this comprehensive approach opens doors for leveraging precision molecular feature analysis and drug repurposing possibilities in KRAS mutant CRC.

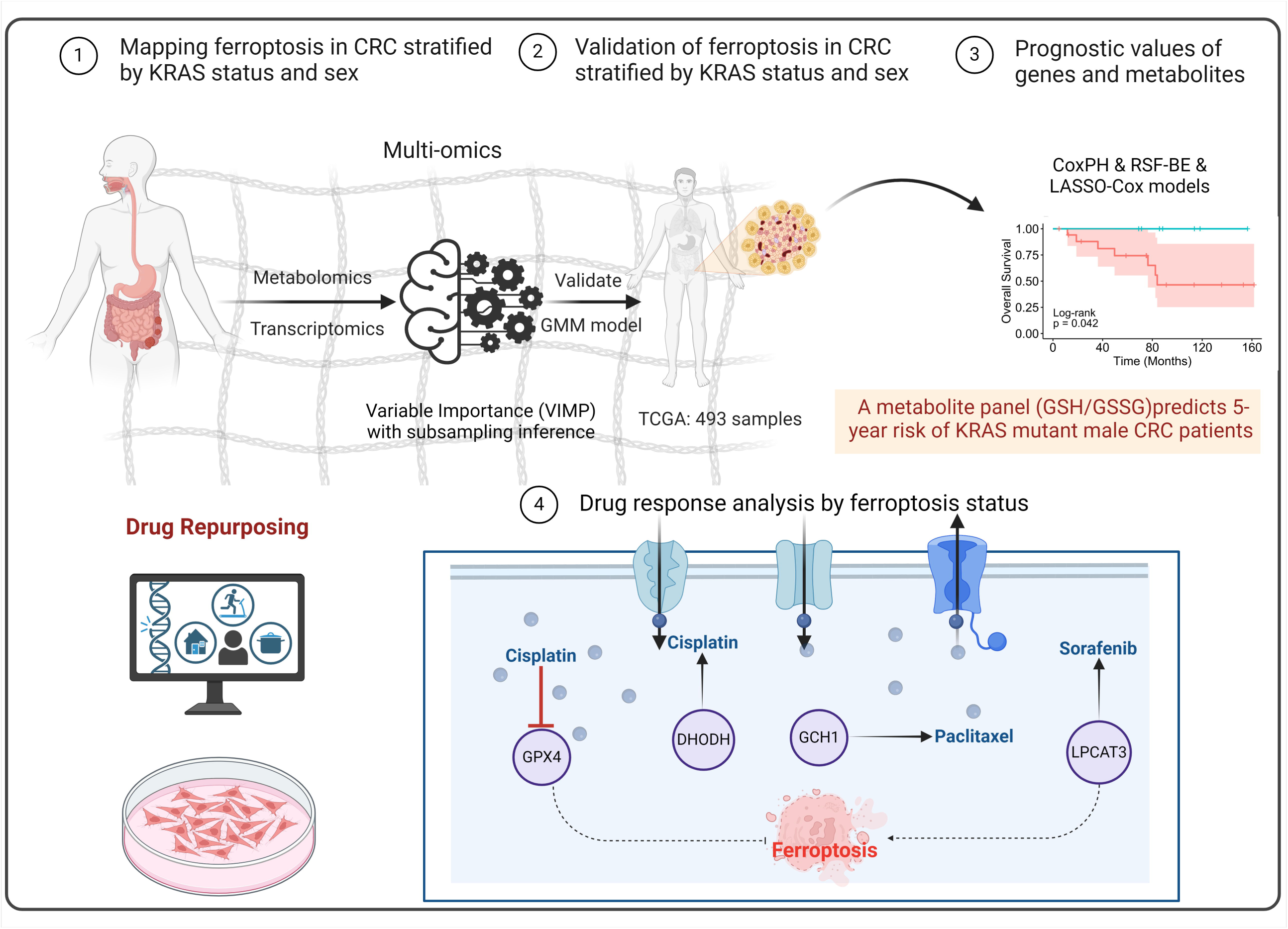

## Introduction

Colorectal cancer (CRC) is characterized by molecular heterogeneity, a high number of mutations, and high mortality(Bray et al., 2018). The KRAS oncogene, is mutated in approximately 40% of metastatic CRC patients, it is both a prognostic and predictive biomarker for which treatment is tailored (Bray *et al*., 2018). Other commonly occurring mutations include BRAF which is present in 7%∼10% in CRCs(Caputo et al., 2019), and TP53 which is present in 43% of all CRC cases (Li et al., 2019). Among these genetic mutations, KRAS has been considered an “undruggable” target in CRC, as KRAS mutations result in resistance to anti-EGFR therapeutics. However, it has become evident that oncogenic KRAS mutations mediate components of the tumor microenvironment, particularly by promoting inflammation and suppressing the immune response, ultimately leading to immune evasion and tumor progression (Hamarsheh et al., 2020; Lal et al., 2018; Liao et al., 2019).

Additionally, CRC incidence and mortality are influenced by the sex of the patient, with global mortality rates in males significantly higher than in females(Brenner et al., 2007). The difference in CRC incidence between females and males is not a new phenomenon and has been observed for at least 25 years(Abancens et al., 2020). Recent evidence also indicates that there is a decreased tendency of KRAS mutant tumors to undergo iron-dependent cell death, or ferroptosis, in male patients only(Yan et al., 2023c). However, the landscape of sex-related differences has not been well characterized in CRCs with KRAS mutations, and remain to be investigated. The signaling networks and the number of heterogeneous features that underlie the mechanisms of sex differences are complex and could potentially impede the development of molecular targeted therapies against KRAS-mutant tumors. Therefore, there is an urgent need to identify reliable sex specific biomarkers which are prognostic and predictive of response to drug therapies in CRC.

Ferroptosis is a non-apoptotic and iron-dependent form of cell death that was discovered in 2012 (Dixon et al., 2012). Ferroptosis is accompanied by oxidation of polyunsaturated fatty acid (PUFA)-containing phospholipids, iron dependency, and the loss of lipid peroxide repair mechanisms(Dixon and Stockwell, 2019). Numerous studies have shown that modulating the ferroptotic cell death pathway can enhance the antitumor effects of drugs for CRC treatment. Similarly, *in vitro* and *in vivo* experiments have revealed that the use of therapeutic agents that trigger ferroptosis-induced cancer cell death (Erastin and Ras-selected lethal 3 (RSL3)) have promise in CRC(Hu et al., 2022; Sui et al., 2018). However, the use of ferroptosis inducers in translational experiments have yet to be initiated, therefore more preclinical work is warranted in future studies. The relationship between ferroptosis and

CRC has been reviewed in detail elsewhere, notably the ferroptotic pathway has numerous molecular features that are prominent in CRC carcinogenesis and thus could be targets for therapeutics (Yan et al., 2023b; Yan et al., 2022). However, there are no large-scale multi-omics studies on ferroptosis in CRC that have examined sex differences. Furthermore, increasing evidence has shown that ferroptosis inhibition or induction can improve drug sensitivity or overcome chemotherapy drug resistance. For example, paclitaxel, is a classical chemotherapy drug that can upregulate protein levels of both p53 and p21 and downregulate solute carrier family 7 member 11 (*SLC7A11)* and *SLC1A5* gene expression in CRC and lung cancer cells; genes that are important in the mechanisms of ferroptosis (Giannakakou et al., 2001; Lv et al., 2017). Bromelain, a dietary supplement used for digestive problems, causes reactive oxygen species (ROS)-induced ferroptosis in KRAS mutant CRC cells via the modulation of Acyl-CoA Synthetase Long Chain Family Member 4 (*ACSL4)*(Park et al., 2018). Furthermore, siramesine combined with lapatinib can induce ferroptosis by increasing ROS levels in breast cancer cells(Ma et al., 2016). Most efforts focused on inducing ferroptosis have been shown to promote cancer therapy efficacy.

In this study, we integrated multi-omics (metabolomic, transcriptomic) data to comprehensively characterize the ferroptosis signaling network and assess biological heterogeneity, and to identify the ferroptosis landscape according to sex and KRAS status in CRC. Bioinformatic machine learning methods (Variable Importance (VIMP) with subsampling inference) and Gaussian mixed models were used to identify genes and metabolites predictive of KRAS status. Random survival forest combined with a backward elimination (RSF-BE) algorithm using1000 bootstraps was utilized to obtain predictive genes and metabolites related to ferroptosis by sex and KRAS mutational status. Furthermore, the associations between ferroptosis gene expression levels, metabolite abundances, and drug responses were investigated. Through this comprehensive landscape of sex differences and KRAS mutations, our study provides biological insight into the therapeutic targeting of ferroptosis.

## STAR★Methods

A schematic depiction of the study design is illustrated in **Figure 1**.

**Figure 1.**
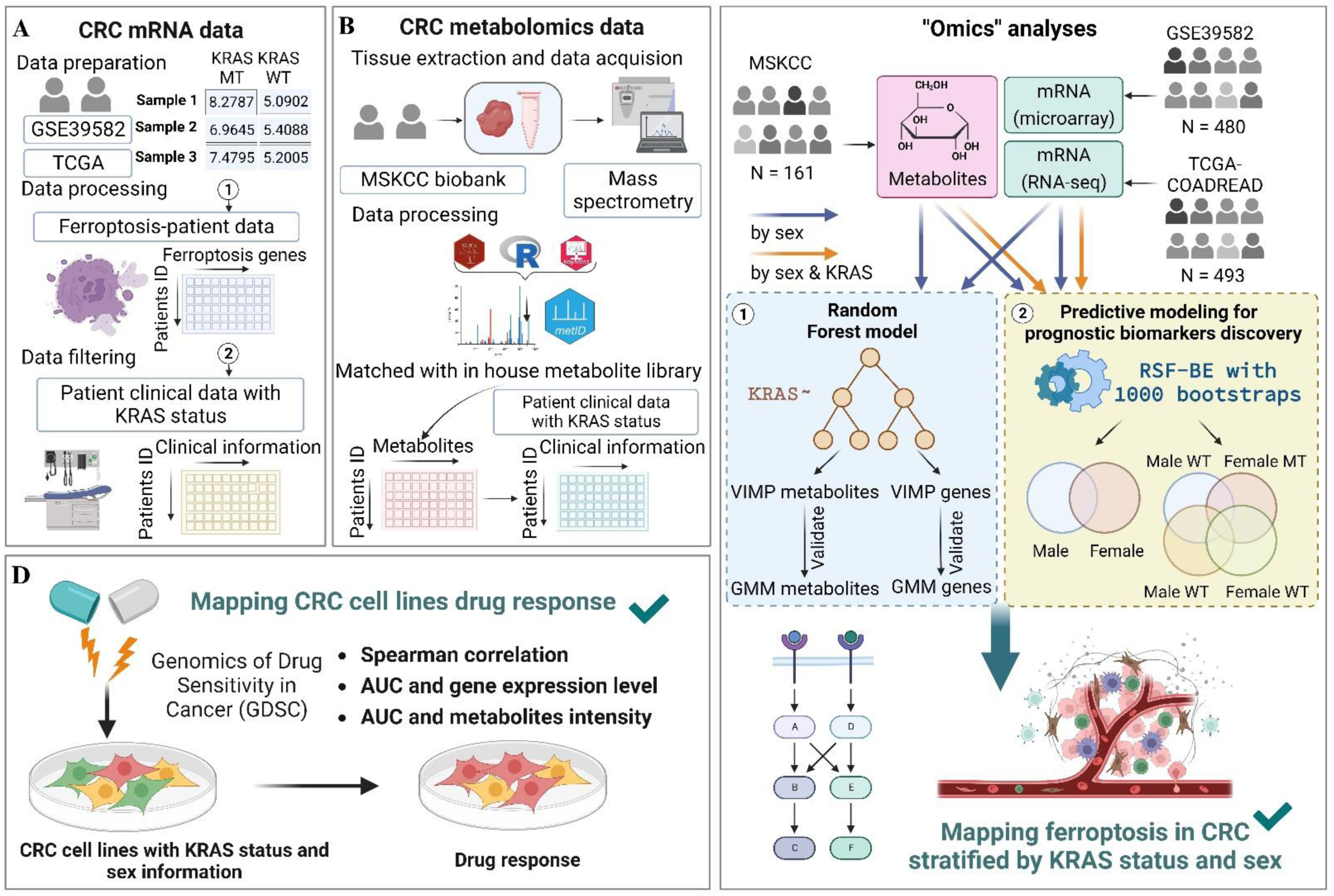
Schematic depiction of the study design. Colorectal cancer (CRC) tissues (stage I-III) were collected from CRC patients. Sex of the patient, KRAS mutation status, gene expression or metabolomics data were collected. Ferroptosis driver and suppressor genes were matched with a gene list from FerrDB. Permutation importance was calculated to assess the importance of each ferroptosis gene by KRAS mutation status. Predictive models using random survival forest with 1000 bootstraps were constructed and predict prognosis by ferroptosis-related gene and metabolite expression. A) Data processing and collection for ferroptosis-related mRNA expression from GSE39582 and TCGA. B) Metabolomics analysis and data processing using tissues from MSKCC cohort. C) Schematic workflow that 1) identified KRAS-dependent genes and metabolites by sex; 2) performed predictive modeling for prognostic biomarkers discovery by sex and KRAS status. D), CRC drug response mapping in silico by analyzing Genomics of Drug Sensitivity in Cancer (GDSC).

### Gene expression data collection and ferroptosis genes collection

Transcriptomics data were collected from three different publicly available patient cohorts.

The 1^st^ gene expression microarray dataset GSE39582 was retrieved from NCBIs gene expression omnibus (GEO) database (https://www.ncbi.nlm.nih.gov/geo/). The dataset contains mRNA expression profiles (Affymetrix U133Plus2, log2 normalized intensity signal) from 585 colorectal tumor tissues and 19 non-tumoral colorectal mucosal tissues. Within this dataset, tumor tissue samples were selected from stage I-III patients (n = 480) across all age; information regarding KRAS type, and sex of patient can be observed in **Table S1**. Relevant to this study, this cohort contains gene expression information on ferroptosis related genes and clinical outcomes and KRAS status information on the 480 patients for 217 females and 263 males with colon adenocarcinoma.

The 2^nd^ transcriptomics dataset used in this study consisted of RNA-seq data (we used log_2_ transformed RSEM (Batch normalized from Illumina HiSeq_RNASeqV2)) and was obtained from two patient cohorts Colorectal Adenocarcinoma TCGA PanCancer Atlas (TCGA-COADREAD) databases which includes both Colon Adenocarcinoma and Rectal Adenocarcinoma downloaded from cBioPortal (https://www.cbioportal.org/study/summary?id=coadread_tcga_pan_can_atlas_2018). We filtered the databases by the same criteria (stage I-III across all ages, information regarding KRAS mutation type, and sex) and obtained data from 237 females and 256 males (**Table S1**). This dataset was used as an independent external parallel dataset tother with GSE39582 to produce consistent findings across platforms and cohorts.

Information on genes that regulate ferroptosis were downloaded from FerrDb which is the world’s first database dedicated to ferroptosis regulators and ferroptosis-disease associations (http://www.zhounan.org/ferrdb/current/#). The database includes ferroptosis driver genes and suppressor genes. We used the ferroptosis genes list to match to genes represented in the two datasets (GSE39582 and TCGA) to produce a ferroptosis gene expression data matrix for VIMP analysis. In TCGA data, 23 genes with severe missing data issues (over 30% missing expression values) were excluded in the modeling and subsequent analyses.

### Metabolomics data, study population and tissue collection

A semi-targeted metabolomics analysis was previously performed on 161 tumor tissues which identified 106 metabolites (Yan *et al*., 2023c). Using clinical records, the patients were assigned to KRAS mutant or KRAS wild type groups, all KRAS mutant subtypes were analyzed together due to sample size. The details of the sample analysis have been published elsewhere, but in brief, the tissues were acquired from surgery on stage I-III CRC patients in the period 1991–2001 at Memorial Sloan-Kettering Cancer Center (MSKCC, New York, NY, United States)(Cai et al., 2020; Yan *et al*., 2023c). The clinical characteristics for the cohort are listed in **Table S1**.

### Variable Importance (VIMP)

The “randomForestSRC” package (version 3.2.2) used in R provides various options for calculating VIMP(Ishwaran et al., 2021). Here, permutation importance was used that adopts a prediction-based approach by using a prediction error attributable to the variable. Permutation importance estimates the prediction error by making use of out-of-bootstrap (OOB) cases. The original data left out from the bootstrap sample used to grow the tree; approximately 1−.632 = .368 of the original sample. This data is called OOB, and prediction error obtained from it is called OOB error. Permutation importance permutes a variable’s OOB data and compares the resulting OOB prediction error to the original OOB prediction error - the motivation being that a large positive value indicates a variable with predictive importance.

We filtered variables (genes and metabolites) with permutation VIMP (P<0.05, signif= “TRUE”). All the significant variables can then be used for survival analysis.

Steps:

1. Train a model f well (assume that the eigenmatrix is X, the target variable is y, and the error measurement index L(y, f)).
2. Through loss function to calculate the original model error ε^orig^ = L (y, f (X)) (MSE)
3. Generate a characteristic field data (for example: mother age) randomly, then throw into the back to before the trained model f calculation error of the calculation model for = L (y, f (X^perm^)).
4. Calculate the factor to the importance of the importance of = ε^perm^ - ε^orig^.
5. Restore the scrambled features in step 4, replace the next feature, and repeat steps 3 to 4 until all features have been calculated.

### High-dimensional Gaussian mixture model

Our goal is to identify a set of genes that are differentially expressed in two groups or a set of genes that would discriminate two groups. Conventional methods such as those that conduct two-group t tests for each individual gene, or Support Vector Machines (SVMs) may fail in this specific problem mainly for two reasons. First, there are correlations among the expression levels of these genes, conducting t tests for individual genes ignores the correlations among the gene expressions. This may lead to spurious detection or failure to detect the genes that truly separate the two groups. Second, SVM involves choosing some kernels that are dependent on unknown quantities. Different choices of kernels may lead to distinct results.

In this paper, we use a robust and efficient method to identify the set of genes that best separate the two groups. We model the expressions of *p* genes of *n* individuals by a Gaussian mixture model (Reynolds, 2009). More details were provided in **Supplementary Document.** MATLAB code is available in Github (https://github.com/yanhongxuejiao/Ferroptosis_CRC).

There are two advantages for using the Gaussian mixture model as follows. First, it is interpretable from a biomedical perspective. The gene expression pattern of each group approximately follows a multi-variate Gaussian distribution because people within the same group have similar but slightly divergent gene expression patterns to account for individual variability. Threfore, the gene expression data when pooling the two groups together would follow a mixture Gaussian distribution with two components, *ω_1_N(μ_1_,Σ)+ ω_2_N(μ_2_,Σ)*with two centers *μ_1,_ μ_2_* well separated(Cai et al., 2019).

The multivariate normality of the gene expression data is checked using the R package MVN (version 3.5.0)(Korkmaz et al., 2014). And hence, it is valid to model the gene expressions by a Gaussian mixture model. When fitting a Gaussian mixture model, we are seeking a “best discriminating direction” to separate two groups. This discriminating direction represents a linear combination of only a small portion of gene expressions. These genes might be involved in some disease-causing pathways and interact with each other, so they together lead to some differential phenotypes between the two groups.

Due to the high-dimensional nature of this problem (*n < p*), we automatically select the set of genes whose expressions best separate two groups after fitting the Gaussian mixture model.

Second, there is well-established statistical methodology to fit a high-dimensional Gaussian mixture model(Cai *et al*., 2019). It is computationally efficient and theoretically guaranteed. The workflow for the entire implementation is carried out using MATLAB equipped with 16GB of RAM. More details are provided in **Supplementary Document.** MATLAB code is available in the Github link (https://github.com/yanhongxuejiao/Ferroptosis_CRC).

### Statistical analysis and variable definitions

Kaplan-Meier (KM) analysis was adopted to measure survival differences between groups (overall survival (OS) and progression-free survival (PFS)) using the R package “survminer” (version 0.4.9). Patients were categorized into early stage (clinical stage I and II) and late stage (clinical stage III) cases. Metabolite abundances in tumor tissues from the MSKCC tumor tissue cohort were log_2_-transformed before modeling. This study used a 2-sided P value < 0.05 as statistical significance level. For multiple comparison, P values were further adjusted using the false discovery rate (FDR).

### Random survival forest with backward elimination (RSF-BE) algorithm

An RSF is computed by a cluster of binary decision trees that have been frequently used to select the most important variables linked with time to event(Wang and Li, 2017). Minimal depth measurement is implemented to assess how informative a variable is regarding the time until event. Harrell’s concordance index (C-index) is equal to 1-prediction error rate, which is commonly applied to evaluate the predictability of a model. We applied a RSF with a tiered backward elimination procedure (BE) reformed based on the RSF-BE method proposed by Dietrich et al.(Dietrich et al., 2016) for variable selection to detect the most predictive and informative features (genes, metabolites) combination while forcing covariates into the model. As **Figure S1** shows, the tiered BE procedure is an algorithm that repeatedly computes RSF using input variables, ranks the features by minimal depth and removes those ranked worst in a dynamic way based on the current feature size, and uses remaining data to compute a new RSF, until there remains one feature, and finally selects a combination of features with minimal error rate (or highest C-index)(Dietrich *et al*., 2016). This dynamic approach ensures both model accuracy and moderate computational burden to allow bootstrapping mentioned in the next section. Models were trained using the “randomForestSRC” package (version 3.2.2) in R software (version 4.3.1).

### Identification of sex-specific predictive genes and metabolites for CRC 5-year OS

In each RSF enforcing inclusion of the two covariates, parameters adopted 300 as ntree to balance model performance and computational burden, nsplit was set as 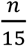 (15 randomly chosen values for each feature), and node size was chosen to be the rounding integer p=of 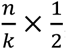, aiming to balance granularity with the risk of overfitting. C-index was used to evaluate the model performance. For GSE39582 and TCGA with mRNA data, RSF-BE was performed on the datasets containing expressions levels of ferroptosis genes plus important prognostic covariates (age and clinical stage) to identify sex-and KRAS-specific predictive signatures for 5-year overall survival (OS) and 5-year progression-free survival (PFS), an outline of the methodology can be seen in (**Figure 2**). The RSF-BE modeling was conducted with 1000 bootstrap resampling iterations for each of the stratified subgroups: 1) males, 2) females, 3) males with KRAS mutant (MT), 4) females with KRAS MT, 5) males with KRAS wild type (WT), and 6) females with KRAS WT. This iterative resampling aimed to generate a distribution of the importance of each feature (gene) to ensure stability and repeatability in the gene selection process. The frequency with which each gene was selected as a predictive feature within the 1000 bootstraps was recorded.

**Figure 2.**
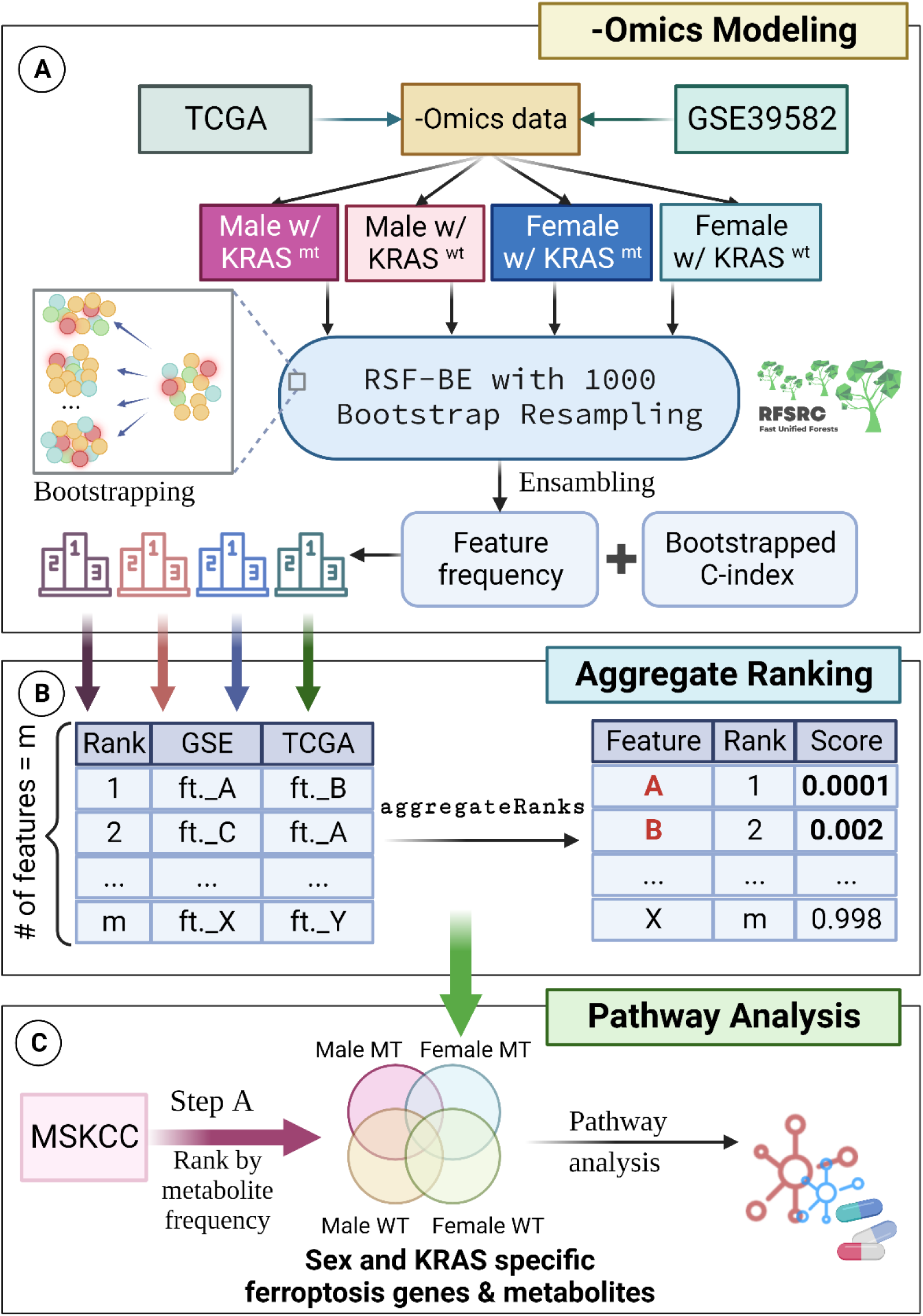
Machine learning-based survival analysis and predictive modeling flowchart. Sex- and KRAS-stratified survival analysis and predictive modeling for each of TCGA, GSE39582, and MSKCC datasets that contained clinical variables and ferroptosis-related-omics data. A) Datasets were stratified by sex and KRAS status. RSF-BE with 1000 bootstraps resampling was applied to the four subgroups (females with KRAS MT, females with KRAS WT, males with KRAS MT, and males with KRAS WT) in GSE39582 and TCGA data. Genes from a certain subgroup were compared based on frequencies of being selected as predictive genes within the 1000 bootstraps between GSE39582 and TCGA cohorts and B) ranked using aggregate ranking method. Genes with scores below 0.05 were selected for each subgroup in further steps. C) MSKCC metabolomics data underwent step A and metabolites were ranked by frequency. Metabolites with frequencies over 300 remained. Therefore, sex and KRAS-specific ferroptosis genes and metabolites were determined and used to perform pathway analysis and enrichment analysis.

Following the RSF-BE modeling, genes from each subgroup were ranked based on their selection frequencies between the GSE39582 and TCGA-COADREAD cohorts. The Robust Rank Aggregation method(Kolde et al., 2012) using R package aggregateRanks (Version 1.2.1) was applied to provide a significant score for each gene across the two datasets where smaller scores indicating higher ranks agreed across different datasets. Genes with aggregate scores below 0.05 were deemed significant and retained for further analyses.

For the MSKCC dataset, a similar RSF-BE methodology was applied. However, instead of genes, the emphasis was on ranking metabolites based on their selection frequency. Metabolites with frequencies exceeding 300 across the bootstrap iterations were selected for subsequent analyses.

Computational modeling process was assisted with the McCleary HPC cluster on Yale Center for Research Computing. Codes are available in https://github.com/XinyiShen0304/Ferr_CRC_Project or https://github.com/yanhongxuejiao/Ferroptosis_CRC.

### Partial dependence plots

Partial dependence survival plots for sex- and KRAS-specific ferroptosis-related prognostic genes and metabolites were plotted to depict non-proportional hazard responses and non-linear relationships between the predicted prognosis and identified genes or metabolites. They were constructed by integrating out the effects of targeted variables beside other covariates and averaging results from the 1000 bootstraps. For plots of metabolites from males with KRAS MT, females with KRAS MT, males with KRAS WT, and females with KRAS WT from MSKCC cohort, we adopted 10 unique data points due to limited sample sizes. For the plots of genes, 25 data points were included, which is the default setting of the plot.variable function from the randomForestSRC package.

### Pathway and Enrichment Analysis

With the selection of sex and KRAS-specific ferroptosis genes and metabolites from the preceding steps, we conducted a comprehensive pathway analysis. This included assessing the biological processes, cellular components, and molecular functions these genes and metabolites were involved in using Gene Ontology (GO) enrichment analysis by R package clusterProfiler (version 4.10.0)(Wu et al., 2021). Additionally, joint-pathway analysis was performed using MetaboAnalyst 5.0(Pang et al., 2022) to identify over-represented biological themes and pathways, providing insights into the mechanistic roles of the selected genes and metabolites in the context of ferroptosis in CRC.

### Analysis of drug response in ferroptosis status

The Area Under the Curve (AUC) data and the gene expression matrix for cancer cell lines were downloaded from *Genomics of Drug Sensitivity in Cancer* (*GDSC*)(Yang et al., 2012) (http://www.cancerrxgene.org/downloads) and Dependency Map (DepMap)(Broad, 2020) (https://depmap.org/portal/). In custom downloads option, we selected “all cell lines”, “Drug sensitivity AUC (Sanger GDSC1)”, “Drug sensitivity AUC (Sanger GDSC2)”, “miRNA” and “Metabolomics”. In total 490 drugs were available to explore in these data (GDSC1: 320, GDSC2:171, **Table S2**). GDSC1 (version 1 of the drug response dataset) and GDSC2 (version 2 of the drug response dataset) are two releases from GDSC database. The cell line information includes KRAS status, sex of patient from whom cell lines were retrieved, and tumor type, and is provided in **Table S3**. To assess the drug response in cancer cell lines, we calculated the Spearman correlation between the AUC and gene expression (Log2) of cancer cell lines from GDSC for drug responsiveness (|Rs| >0.5; FDR < 0.05). Additionally, we calculated the Spearman correlation between the AUC and metabolites abundance (Log2) of cancer cell lines from GDSC for drug responsiveness (|Rs| >0.5; FDR < 0.05). As previously reported, a positive Spearman correlation is defined as drug-resistant, while a negative Spearman correlation is defined as drug-sensitive(Ye et al., 2019).

## Results

### Sex-specific VIMP of ferroptosis-related genes and metabolites

We conducted a VIMP analysis to examine the different effects of nodule characteristics on the model performance (**Figure 1C)**. Unlike Gini importance that gauges significance through in-sample impurity, permutation importance uses a predictive model by employing a prediction error linked to the variable. A clever feature is that instead of using cross-validation, which is computationally costly for forests, permutation importance estimates prediction error by utilizing out-of-bootstrap instances(Ishwaran et al., 2022; Louppe et al., 2013). Employing a permutation predictor importance carries three advantages. Firstly, its computation isn’t dependent on a specific model form. Secondly, the predictive model required only a single training. Lastly, the random shuffling can be executed multiple times to decrease variability in the calculation(Ishwaran *et al*., 2022).

As shown in **Figure 3A**, for males represented in the GSE39582 cohort, ferroptosis suppressors *SLC7A11, PPP1R13L, NR4A1, NFS1, GPX4, FTL, FTH1, FH,* and *ACSL3* were significantly different in their expression (- log10P>1.31) between KRAS mutant and KRAS wild type tumors, as were ferroptosis drivers *SLC1A5, ELOVL5, ALOX5, ALOX12B, ALOX12,* and *ACSL4*. However, these differences were not observed in females (- log10P>1.31) for both suppressor and driver genes. In addition, 20 metabolites from the MSKCC cohort were found to be significantly different between KRAS mutant and wild type tumors in male patients, including lactate, lipoxin B4, prostaglandin F2α, stearic acid, histidine, xanthine, methionine, vitamin E and glutamine, while in female patients, only stearic acid was altered and identified to be related to ferroptosis. Those that have sex-differences are shown in **Figure 3B**.

**Figure 3.**
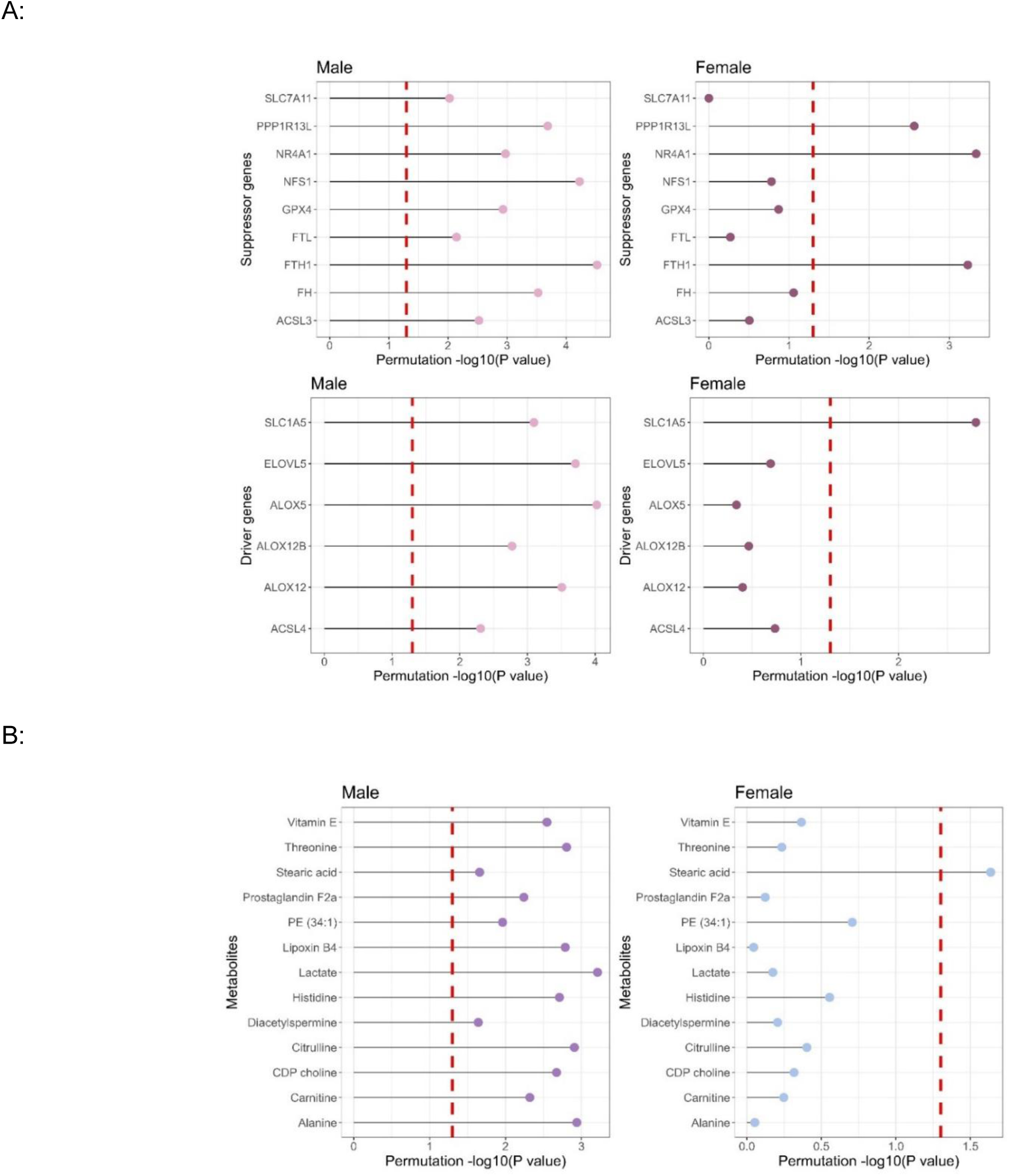
Ferroptosis genes and metabolites in colorectal cancer tissues revealed by VIMP analysis. Permutation P value assessment showing variance of genes between KRAS mutant CRCs and KRAS wild-type CRCs. The results are with respect to KRAS mutant CRCs when compared to wild type CRCs. (A) Genes from GSE39582 Colon tumor tissues from male patients: KRAS mutant tumors (n = 97), KRAS wild type tumors (n = 166); Colon tumor tissues from female patients: KRAS mutant tumors (n = 86), KRAS wild type tumors (n = 131), (B) Metabolites from MSKCC cohort. Colon tumor tissues from male patients: KRAS mutant tumors (n=25), KRAS wild type tumors (n = 58); Colon tumor tissues from female patients: KRAS mutant tumors (n = 35), KRAS wild type tumors (n = 43).

We further examined which genes were overlapped when comparing KRAS mutant and KRAS wild type tumors in the GSE39582 and TCGA-COADREAD datasets. A gene with a positive lower bound of VIMP estimate was considered significantly dependent on KRAS mutation status. As illustrated in **Table S4**, for male patients, 56 genes, including *ALOX12, ALOX5, FTH1, SLC1A5, SLC38A1, SLC3A2*, and *HMOX1*, were retained in both datasets. However, for females, *ALOXs* were not selected in either dataset. Among the 32 overlapping genes, *FTH1, HMOX1* were preserved.

### Sex-specific Gaussian Mixed model of ferroptosis-related genes

Gaussian mixed model analysis was then performed with the same dataset, GSE39582, to validate the results observed from the VIMP analysis. As shown in **Table S5**, in males from the GSE39582 cohort, *TLR4, DPP4, G6PD, FTH1, GPX4, SLC7A11* and others were selected to classify the KRAS status difference which were overlap with VIMP results. However, among all the significant ferroptosis genes in both GMM and VIMP, only *FTH1* was selected in female CRCs (**Table S5**).

### Sex-specific prognostic values of ferroptosis-related genes

The workflow in **Figure 2** shows our method to conduct a deeper exploration of the complex relationship between ferroptosis-related genes, metabolites, and patient prognosis. Our findings underscore the roles that both sex and KRAS mutation status play in regulating the prognostic landscape of CRC. All RSF-BE models, categorized by sex and KRAS status across the two cohorts (GSE39582 and TCGA-COADREAD), demonstrated high efficacy with an average bootstrapping C-index nearing or exceeding 0.9 (**Figure S2**). RSF-BE models applied to GSE39582, TCGA, and MSKCC cohorts all yielded discriminable genes and metabolites by sex and KRAS status based on their ranks with clear gradients ranging from 0 to above 900 within the 1000 iterations (**Table S6**). The rankings of genes from each subgroup for GSE39582 and TCGA were then calculated by Robust Rank Aggregation(Kolde *et al*., 2012). With a cut-off of 0.05 for the score value, unique and overlapping genes were finally determined that are highly predictive of 5-year OS in both datasets when stratified by sex and KRAS status (**Figure 4A, Table S7, Figure S3)**. Here, it is evident that different sets of genes play pivotal roles in females and males, further delineated by KRAS mutation status (MT and WT). Similarly, sets of metabolites (**Figure 4B**) present distinctions between sex and KRAS status among prognostic metabolites (**Table S6**). With a closer look at males with KRAS mutations who were reported to have decreased ferroptosis(Yan *et al*., 2023c), partial dependence plots (**Figure 4C**) elucidated the non-linear relationships between predicted OS probabilities and 15 selected ferroptosis genes (*AIFM2*, *AQP8*, *CDCA3*, *CYGB*, *GCLC*, *HMOX1*, *IDO1*, *INTS2*, *KDM6B*, *MAP3K14*, *MEG3*, *PEX6*). For instance, *CYGB*, *GCLC*, and *HMOX1* had inverted U shaped non-monotonic trends of predicted 5-year and 3-year OS, with increasing, followed by decreasing OS probabilities. **Figure S4** displays the remaining partial dependence plots for the other subgroups using GSE39582 cohort. Likewise, the non-linear and non-monotonic relationships between metabolite abundance and predicted overall survival for the MSKCC cohorts are depicted in **Figure S5**.

**Figure 4.**
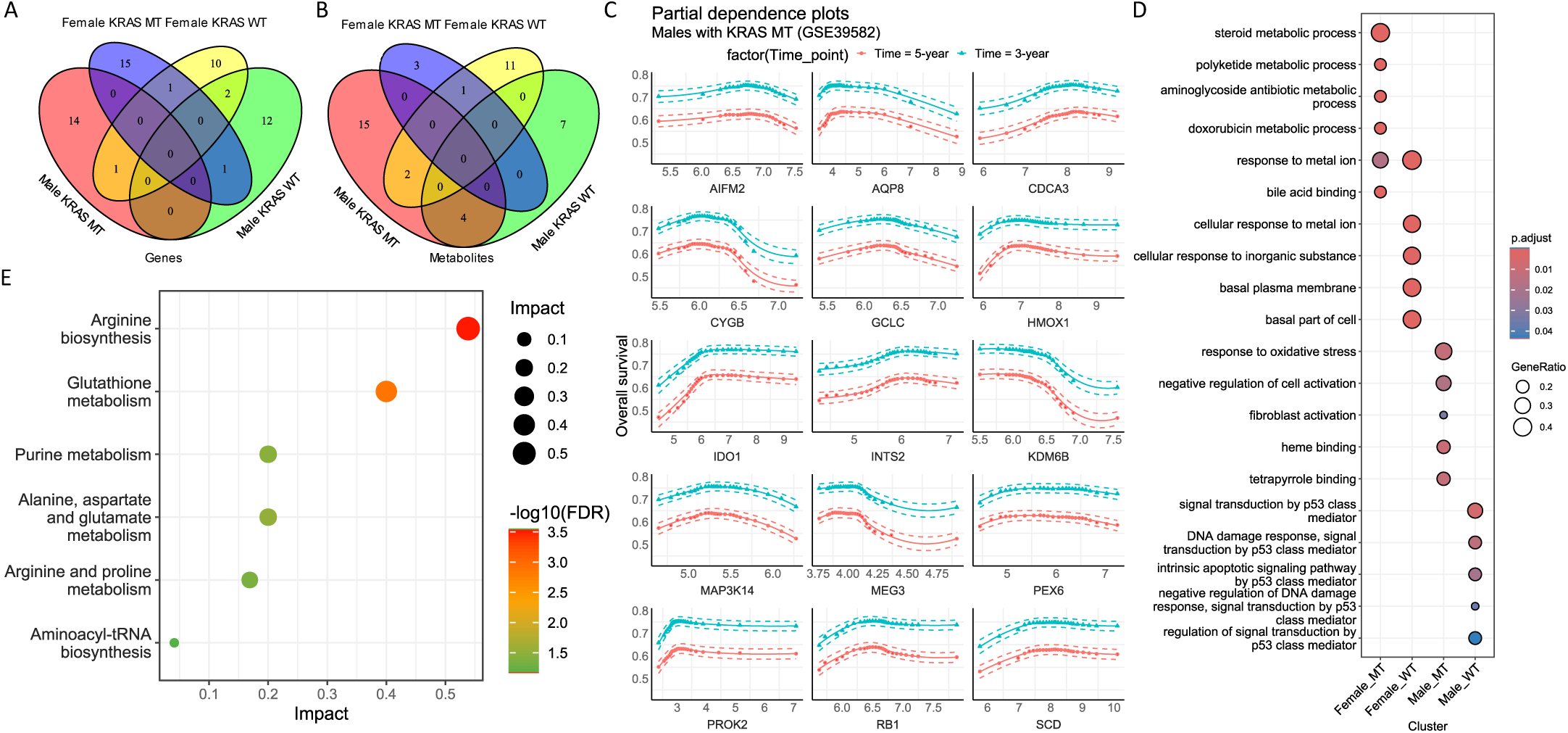
Ferroptosis-related genes and metabolites are predictive of prognosis in a sex- and KRAS-dependent manner. A) Distinct sets of ferroptosis genes by sex and KRAS status are predictive of 5-year OS and B) metabolites by sex and KRAS status are predictive of OS. C) Partial dependence plots display non-linear relationships between predicted 5-year and 3-year OS probability with prognostic ferroptosis genes for males with KRAS mutations in the GSE39582 cohort. D) Gene Ontology (GO) enrichment analysis using prognostic ferroptosis genes reveals biological processes, cellular components, metabolic pathways by sex and KRAS status. E) Joint-pathway analysis using prognostic ferroptosis genes and metabolites suggests that arginine biosynthesis and glutathione metabolism were enriched in males with KRAS mutations.

Gene Ontology (GO) enrichment analysis (**Figure 4D**) offered a comprehensive view into the broader biological landscape influenced by these prognostic ferroptosis genes. From steroid metabolic processes to response to oxidative stress and heme binding, the range of biological processes, cellular components, and metabolic pathways shown highlights the potential interactions between ferroptosis genes that would impact prognosis. The significance values from stratified analysis showed varied enrichment levels based on sex and KRAS status.

The joint-pathway analysis (**Figure 4E, Table S8**) integrated the results from genes and metabolites acquired across all three cohorts, offering a holistic view of the prognostic landscape. Among the highlighted pathways, arginine biosynthesis and glutathione (GSH) metabolism stood out with both significant adjusted P values (both Holm’s method and FDR correction) and high pathway impacts, particularly for males with KRAS mutations compared to all the other subgroups with either insignificant adjusted P values or low pathway impacts (**Table S8**).

For a more detailed examination of the impact of sex and KRAS on glutathione metabolism, we classified patients based on the median values of the reduced to oxidized glutathione ratio (GSH/GSSG ratio) from the MSKCC cohort and conducted a Kaplan-Meier analysis. **Figure 5** shows that a lower ratio of GSH/GSSG was linked to worse OS in only males with KRAS mutation, which was not observed in any of the other sex and KRAS mutation subtypes by KM analysis.

**Figure 5.**
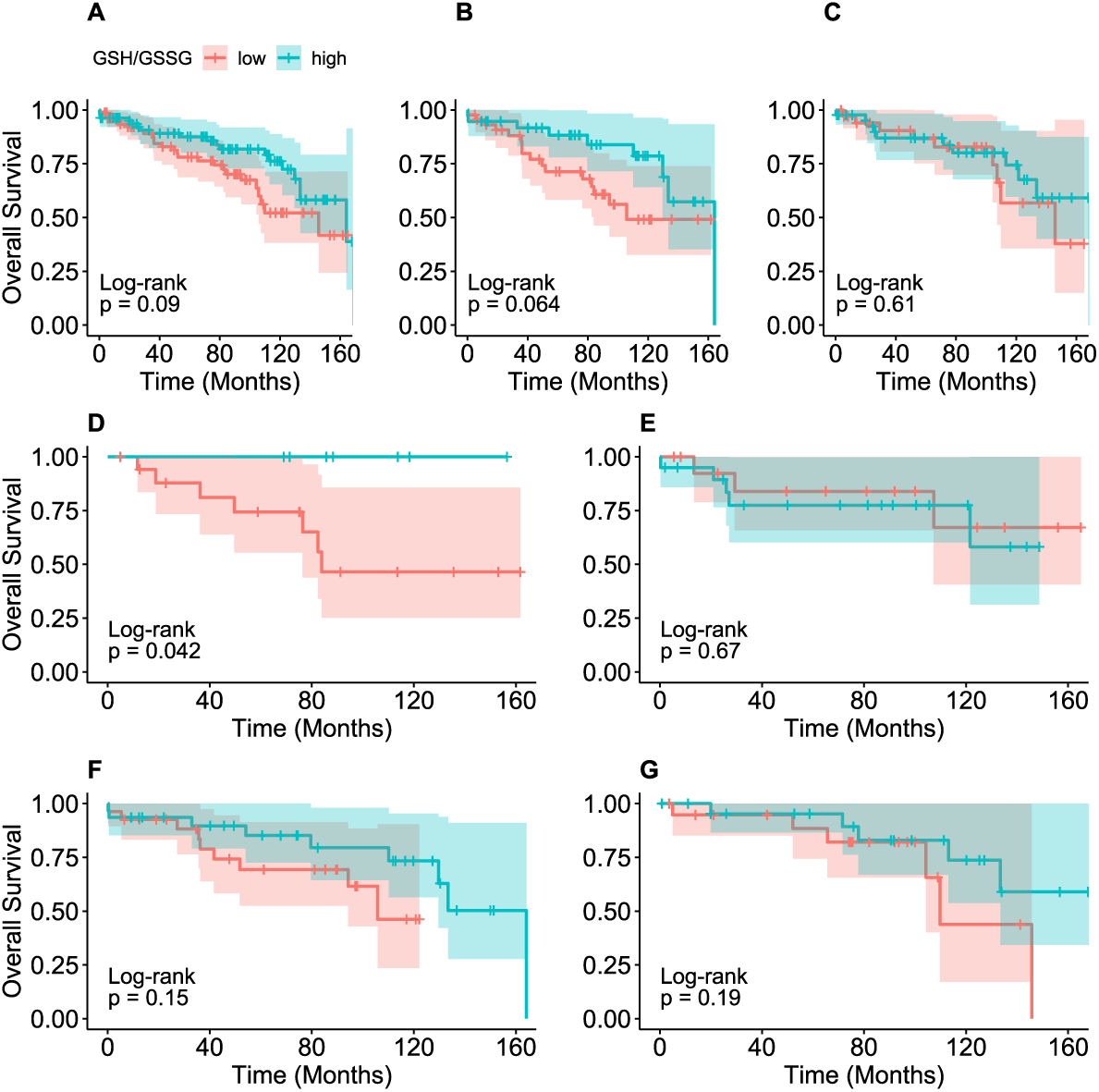
Kaplan-Meier analysis of GSH/GSSH levels stratified by sex and KRAS status using metabolomics data from the MSKCC cohort. A) All patients, B) male patients only, C) female patients only, D) male patients with KRAS MT tumors, E) female patients with KRAS MT tumors, F) male patients with KRAS WT tumors, and G) female patients with KRAS WT tumors.

### Associations between ferroptosis gene mRNA expression and drug response

To investigate the associations between ferroptosis genes (by mRNA expression) and drug response, we examined data obtained from 30 different CRC primary tumor cell lines, and integrated information regarding their KRAS status and sex of the patient from which the cell line originated. Spearman correlations were performed to assess the drug response status across all the cell lines. Within the mechanisms of ferroptosis there are four major surveillance systems. One is the GPX4-GSH system, which reduces phospholipid peroxides (PLOOHs) to corresponding PLs, this results in the inhibition of GSH synthesis which suppresses ferroptosis(Dixon *et al*., 2012; Yang et al., 2014), others are mediated by enzymes such as DHODH, FSP1 and GTP cyclohydrolase-1 (GCH1) that deal with antioxidant activity, thus suppress PL peroxidation. *GPX4* expression in cell lines that originated from male patients with KRAS mutant CRCs was associated with changes in sensitivity to 61 anticancer drugs (Rs<-0.5, FDR<0.05, **Table S9**). However, in KRAS wild type male CRC cell lines *GPX4* expression was associated with changes in sensitivity to just 37 drugs. Upon examination of the overlap between KRAS mutant and wild type cells for *GPX4* and drug sensitivity, there were just two drugs (Rucaparib and SB216763), whereas *GPX4* expression had no correlations to drug sensitivity in cell lines from female CRC patients. However, *DHODH* expression in cells from male KRAS mutant CRCs was associated with resistance to 98 drugs (Rs>0.5, FDR<0.05, **Table S9**), conversely, in cell lines from female KRAS mutant CRCs, *DHODH* was associated with sensitivity of 24 drugs. In addition, *AIMF2 (FSP1)* was associated with resistance to 97 drugs in cells from male KRAS mutant CRC but associated with sensitivity of 19 drugs in cells from female KRAS mutant CRCs. Furthermore, *GCH1* expression in cells from male KRAS mutant CRCs was associated with sensitivity of mirin and resistance of ponatinib, paclitaxel, PD173074. In cells from female KRAS mutant CRCs, *GCH1* was associated with sensitivity of 25 drugs (Rs<-0.5, FDR<0.05, **Table S9**).

For other ferroptosis driver genes, we found that *ACSL4* in cells from male KRAS mutant CRCs was associated with sensitivity of 15 anticancer drugs (Rs<-0.5, FDR<0.05, **Table S9**) which included mTOR, PI3K, and WNT signaling target drugs, but this sensitivity was not observed in cell lines from the other three subgroups (male KRAS wild type CRCs, female KRAS mutant CRCs, female KRAS wild type CRCs). However, another gene involved in lipid metabolism *LPCAT3* was associated with resistance to 53 anticancer drugs (Rs>0.5, FDR<0.05, **Table S9**) in cell lines from male KRAS mutant CRCs only. *CBS*, which is involved in transsulfuration activity, was associated with changes to sensitivity in 19 drugs (Rs<-0.5, FDR<0.05, **Table S9**), and *FTL* that regulates iron metabolism was associated with changes in sensitivity of 16 drugs in cell lines from male KRAS mutant CRCs (Rs<-0.5, FDR<0.05, **Table S9**), but not observed in cell lines from the other three subgroups. Furthermore, in arachidonic acid (AA) metabolism, *ALOX12* was associated with resistance to 66 drugs (Rs>0.5, FDR<0.05, **Table S9**), whereas in cell lines from male KRAS wild type CRCs, 18 drugs were associated with *ALOX12* resistance, however, this funding was not observed in cell lines from females. *ALOX15* in cells from male KRAS mutant CRCs was associated with resistance to six drugs (Rs>0.5, FDR<0.05, **Table S9**), additionally, *GLS2*, which has been shown to promote ferroptosis, was associated with resistance to seven drugs and with changes in sensitivity of two drugs (SB505124, tamoxifen) for this subgroup. Regarding classical anti-tumor drugs and compounds associated with ferroptosis in CRC (**Table 1**), several crucial ferroptosis suppressor genes such as *AIFM2* (*FSP1*), *DHODH*, and *FH* exhibit resistance to Cisplatin, which targets GSH-GPXs to induce ferroptosis (**Figure 6**). Additionally, *GCH1* displays resistance to Paclitaxel, which targets SLC7A11, whereas *LPCAT3*, a ferroptosis driver gene, exhibits resistance to Sorafenib, a targeted drug in CRC (**Figure 6**). However, as the backbone drugs to CRC treatment, oxaliplatin and irinotecan were not associated with ferroptosis suppressor and driver genes (FDR>0.05).

**Figure 6.**
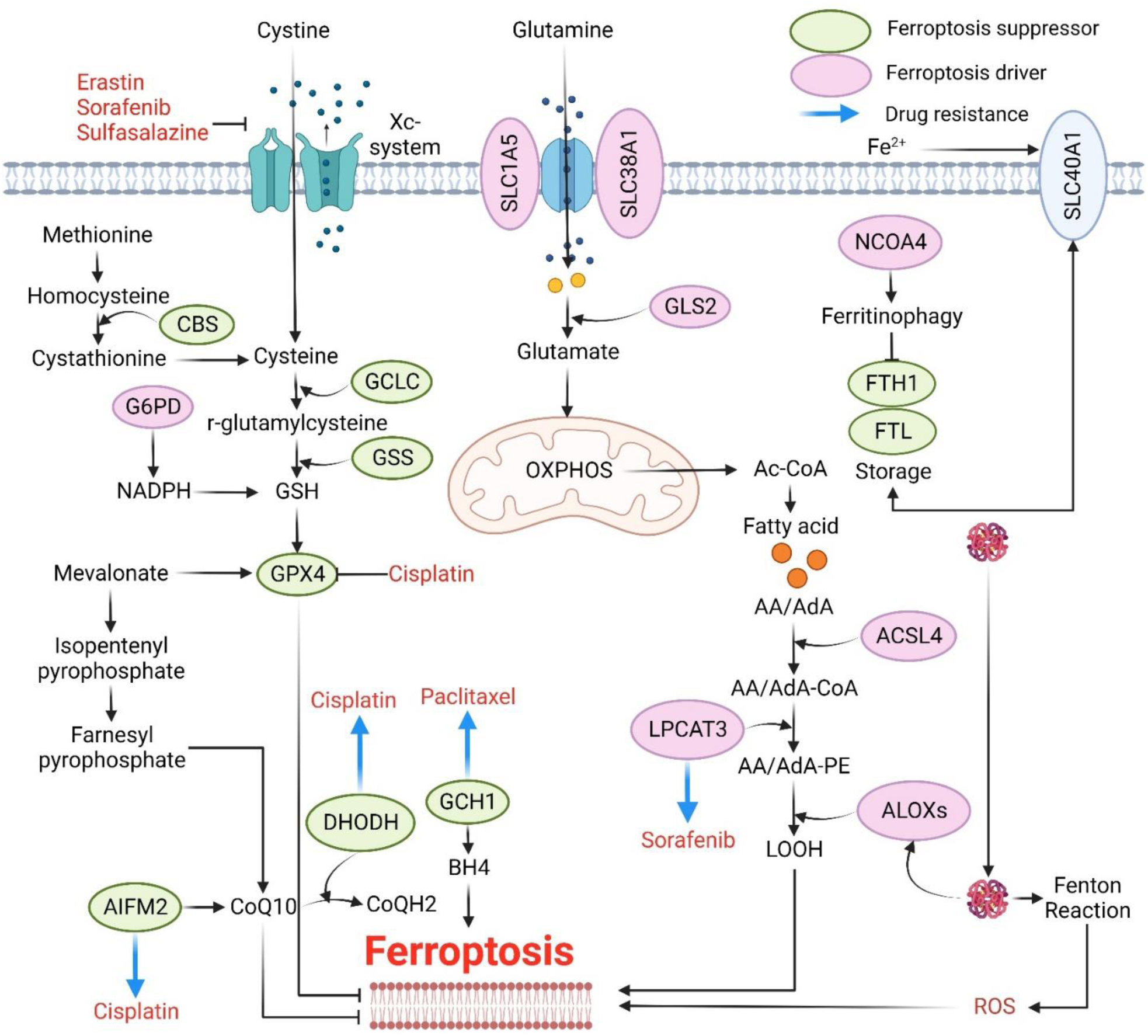
Core molecular machinery and signaling regulation of ferroptosis. Ferroptosis can occur through two major pathways: the extrinsic or transporter-dependent pathway (e.g., decreased cysteine or glutamine uptake and increased iron uptake), and the intrinsic or enzyme regulated pathway (e.g., the inhibition of GPX4). Blue arrows represent resistance to drug, pink genes represent ferroptosis drivers, green genes represent ferroptosis suppressors. Only significant drug response in male KRAS mutant types was presented. Abbreviations: G6PD: glucose-6-phosphate dehydrogenase; GSH: Glutathione; CoQ10: Coenzyme Q10; CoQH2: Ubiquinol; BH4: Tetrahydrobiopterin; AA: Arachidonic acid; AdA: Adrenic acid; CoA: Coenzyme A; PE: Phosphatidylethanolamine; OXPHOS: Oxidative phosphorylation; CBS: Cystathionine-Beta-Synthase; GCLC: Glutamate-Cysteine Ligase Catalytic Subunit; GSS: Glutathione Synthetase; GPX4: Glutathione Peroxidase 4; NADPH: Nicotinamide adenine dinucleotide phosphate; AIFM2: Apoptosis Inducing Factor Mitochondria Associated 2; DHODH: Dihydroorotate Dehydrogenase; GCH1: GTP Cyclohydrolase 1; LPCAT3: Lysophosphatidylcholine Acyltransferase 3; ALOX: Arachidonate 5-Lipoxygenase; ACSL4: Acyl-CoA Synthetase Long Chain Family Member 4; FTL: Ferritin Light Chain; FTH1: Ferritin Heavy Chain 1; NCOA4: Nuclear Receptor Coactivator 4; GLS2: Glutaminase 2; SLC1A5: Solute Carrier Family 1 Member 5; SLC38A1: Solute Carrier Family 38 Member 1; SLC40A1: Solute Carrier Family 40 Member 1

**Table 1.**
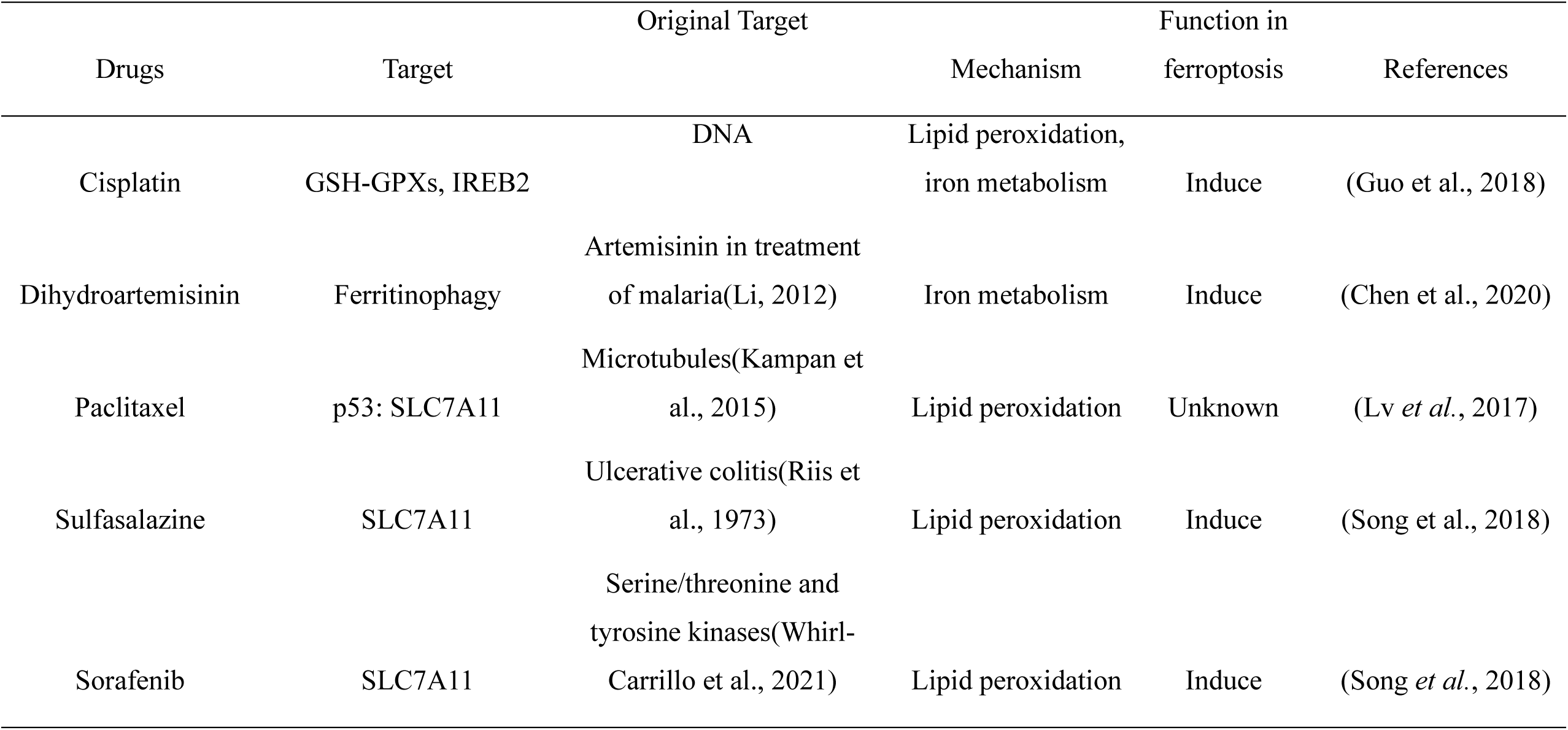
Classical anti-tumor drugs and compounds associated with ferroptosis in CRC.

| Drugs | Target | Original Target | Mechanism | Function in ferroptosis | References |
| --- | --- | --- | --- | --- | --- |
| Cisplatin | GSH-GPXs, IREB2 | DNA | Lipid peroxidation, iron metabolism | Induce | (Guo et al., 2018) |
| Dihydroartemisinin | Ferritinophagy | Artemisinin in treatment of malaria(Li, 2012) | Iron metabolism | Induce | (Chen et al., 2020) |
| Paclitaxel | p53: SLC7A11 | Microtubules(Kampan et al., 2015) | Lipid peroxidation | Unknown | (Lv *et al.*, 2017) |
| Sulfasalazine | SLC7A11 | Ulcerative colitis(Riis et al., 1973) | Lipid peroxidation | Induce | (Song et al., 2018) |
| Sorafenib | SLC7A11 | Serine/threonine and tyrosine kinases(Whirl-Carrillo et al., 2021) | Lipid peroxidation | Induce | (Song *et al.*, 2018) |

### Associations between ferroptosis metabolites and drug response

Similarly, we investigated the associations between metabolites and drug responses, again focusing on the 30 CRC primary tumor cell lines and the role of KRAS status and sex. According to the mechanisms governing ferroptosis, there are several main metabolic pathways, namely, the transsulfuration pathway, GSH biosynthesis, AA metabolism, the tetrahydrobiopterin (BH4) cycle, and the mevalonate pathway(Yan et al., 2023a; Yan et al., 2023e). In the transsulfuration pathway, S-adenosylhomocysteine (SAH) is associated with changes to the sensitivity of eight anticancer drugs (Rs<-0.5, FDR<0.05, **Table S10**) in cell lines from male KRAS mutant CRCs but was associated with changes to resistance of 17 drugs (Rs>0.5, FDR<0.05, **Table S10**). No significant associations were observed in cell lines from female patients. Another metabolite, methionine, was also associated with resistance to 84 drugs (Rs>0.5, FDR<0.05, **Table S10**). In addition, homocysteine was associated with changes to sensitivity in 15 drugs ((Rs<-0.5, FDR<0.05, **Table S10**) in cell lines from KRAS mutant CRCs, while significant drugs are resistant in KRAS wild type CRCs. In GSH metabolism, cystathionine, which is the upstream metabolite of cysteine, was associated with changes to sensitivity of nine drugs (Rs<-0.5, FDR<0.05, **Table S10**) in cell lines from male KRAS mutant CRCs. GSH was associated with changes in resistance to 52 drugs (Rs>0.5, FDR<0.05, **Table S10**) in cell lines from male KRAS mutant CRCs, including sorafenib (**Table 1**). Oxidized glutathione (GSSG) was associated with changes to sensitivity of seven drugs and resistance with four drugs in cell lines from male KRAS mutant CRCs, but not in female cell lines.

## Discussion

Ferroptosis is regulated by tumorigenesis and is associated with the cellular response to immunotherapy, hormone therapy, and targeted therapies(Sun et al., 2021). A comprehensive analysis of multi-omics data by KRAS status and sex, and their links to therapeutic responses in ferroptosis-related pathways will contribute to understanding patient-level responses to CRC therapy, especially for patients that harbor KRAS mutant CRC tumors. The complexity of the signaling network, the heterogeneous features, and the interplay with sex-related mechanisms have contributed to the difficulty in developing molecular targeted therapies against KRAS-mutant tumors. Despite the critical roles of ferroptosis in CRC(Yan *et al*., 2023a; Yan *et al*., 2022), there are no applicable methods for estimating ferroptosis genes or metabolites stratified by KRAS status and sex in a large number of patients, and furthermore, the landscape of sex-related difference has not been well characterized in CRCs with KRAS mutations. Our aim was to enhance the understanding of ferroptosis’s impact on KRAS mutant CRC with a focus on sex-specific differences. Furthermore, our investigation into the prognostic relevance of ferroptosis-related genes and metabolites in CRC, stratified by sex and KRAS status, offers vital insights into the multifaceted nature of tumor biology.

In this study, we applied VIMP analysis and Gaussian mixed models to examine multi-omics data obtained from CRC patients with a focus on ferroptosis-specific genes and metabolites. The RSF-BE algorithm was used to identify the predictive combination of biomarkers for CRC prognosis that were unique to sex and KRAS status. Utilizing RSF-BE with 1000 bootstrap resampling, we ensured that a robust methodological approach was used to select features that were not only predictive but also consistently repeated across resampling iterations, minimizing the risk of overfitting and random noise. The partial dependence plots based on our machine learning-based survival analysis added additional understanding of the non-linear relationships between prognostic genes and patient outcomes. Additionally, the integration of genetic and metabolic data through joint-pathway analysis provides a integrative view of the sex and KRAS specific ferroptotic pathways. By integrating results from TCGA, GSE39582, and MSKCC datasets using aggregate ranking techniques, we ensured the findings were generalizable and robust across different cohorts. The enrichment of pathways like arginine biosynthesis and GSH metabolism in males with KRAS mutations underscores potential therapeutic targets.

Analysis of the TCGA dataset revealed potential novel therapeutic targets for CRC(Yan *et al*., 2023b): *SLC7A11, GPX4, ACSL4* which were all significantly different in their expression levels between KRAS mutant and KRAS wild type CRCs from male patients. Ferroptosis suppressors *PPP1R13L, NFS1, FTL, FTH1* and *FH* were also discovered as significantly altered by KRAS genotype in males (**Figure 2**). In our previous study(Yan *et al*., 2023c), we analyzed specific genes related to iron accumulation, iron transport, lipid peroxidation, GSH biosynthesis, transsulfuration pathway, and mevalonate pathway. However, in this study, we performed ferroptosis mapping using GSE39582 and TCGA cohorts, incorporating all known ferroptosis drivers and suppressors from the FerrDB database. We matched our datasets with the gene list from FerrDB and used VIMP and Gaussian mixed model to identify significant variables (genes or metabolites) between KRAS mutant and KRAS wild type CRCs based on sex. This approach enables a deeper investigation of the functional roles of ferroptosis in patients based on their KRAS status and sex.

With regards to the significance of individual molecules in the biology of ferroptosis and cancer, it is known that the regulatory subunit 13-like of protein phosphatase 1 (PPP1R13L) has been discovered to inhibit ferroptosis in lung cancer. This process is contingent on the pathway involving NFE2L2, hypoxia-inducible factor 1 subunit alpha (HIF1α), and transferrin (TF)(Ajoolabady et al., 2021). In lung epithelial type 2 cells, and in mice subjected to intestinal ischemia and reperfusion -induced acute lung injury (ALI), the therapeutic potential of activating NFE2L2 to halt ferroptosis by restoring cellular redox balance was demonstrated(Ajoolabady *et al*., 2021; Li et al., 2020). Additionally, NFS1 activity can maintain iron-sulfur co-factors present in cell-essential proteins upon exposure to oxygen compared to other forms of oxidative damage(Alvarez et al., 2017). In our previous study, NFS1 was only upregulated in KRAS mutant tumors compared to normal tissues in males that indicated suppressed ferroptosis in tumors from KRAS mutant male patients(Yan et al., 2023d). FH, also known as fumarase, is a metabolic enzyme in TCA cycle, and a mitochondrial tumor suppressor(Alam et al., 2005). Recent evidence showed that enhanced ferroptosis can promote the suppression of tumors by FH(Gao et al., 2019).

For iron metabolism, ferritin light chain (FTL) or ferritin heavy chain (FTH1)(Muhoberac and Vidal, 2019) plays a pivotal role as the main regulator. FTH1, specifically, is responsible for regulating ferritin activity by controlling the oxidation of Fe^2+^ to Fe^3+^. Notably, in cells with RAS mutations, decreased ferritin expression contributes to heightened sensitivity to ferroptosis, as it leads to increased iron uptake and reduced iron storage(Kim et al., 2021; Yang and Stockwell, 2008). In our study, *FTH1* and *FTL* are significant in males, but not in females. Furthermore, the RSF-BE analysis showed that higher expression of *FTH1* was linked to a worse 5-year OS which is consistent with our previous study(Yan *et al*., 2023d).

Our analysis further suggested that *ALOX* genes are potential targets to mediate ferroptosis induction and sensitize tumor cells. Consistent with our previous findings(Yan *et al*., 2023c), tumors from KRAS mutant male patients had altered AA metabolism (in comparison to KRAS wild type), *ELOVL5, ALOX5, ALOX12B* and *ALOX12* were also significantly altered, but no differences were seen between the genotypes in tumors from female patients. Of note, *ELOVL5* was previously shown to maintain intracellular levels of AA and adrenic acid in gastric cancer, together with regulation by fatty acid desaturase 1 (FADS1), thus could be a potential checkpoint in the ferroptosis pathway(Lee et al., 2020).

ALOXs are iron-containing dioxygenases that insert oxygen into PUFAs in a stereospecific manner(Kuhn et al., 2005). The role of ALOX in ferroptosis depends on the cellular environment due to differences in tissue and cell expression profiles among ALOX family members. For example, only *ALOX5*, *ALOX12*, or *ALOX15* overexpression sensitizes human embryonic kidney cells to ferroptosis(Shah et al., 2018).

Importantly, we have revealed a unique relationship between the GSH metabolic pathway and CRC prognosis dependent on KRAS status, and sex. In MSKCC cohort, tumors from KRAS mutant males with low GSH/GSSG ratios exhibited a worse OS. An alteration in the GSH/GSSG ratio could disrupt cellular homeostasis, wherein GSH levels decreased and GSSG levels increased, which indicated that the activity of GPX4 was compromised. This might lead to an accumulation of lipid peroxides that triggered ferroptosis. In males with KRAS mutations, the significant difference in OS by GSH/GSSG ratio might be attributable to the link between GSH metabolism and CRC progression. It has been shown that the GSH/GSSG ratio in serum from CRC patients was lower in cases than healthy controls and was increased in CRC patients after they received treatment(Acevedo-Leon et al., 2021). However, recent literature has hinted at the differential response of males and females to oxidative stress, an imbalance that is known to promote carcinogenesis and alter cancer progression(Allegra et al., 2023; Wang et al., 2020), providing insights into the role of sex-specific differences in oxidative stress responses and their subsequent impact on cancer susceptibility. These further indicated that the inherent biological sex differences may influence their response to GSH metabolism imbalance. Arginine synthesis, which was enriched in KRAS mutant male tumors, is involved in the biosynthesis of polyamines. Even though there is limited direct evidence between arginine and the regulation of ferroptosis, polyamines have been reported to regulate ferroptosis, therefore arginine as a precursor of polyamine could play a role(Ou et al., 2016; Stewart et al., 2018). Recent evidence suggested that deprivation of arginine effectively hindered erastin-induced ferroptosis, while without impact on RSL3-induced ferroptosis in various types of mammalian cells(Guo et al., 2023). We therefore hypothesize that ferroptosis, redox imbalance, and genetic mutations including KRAS might not only determine the course of CRC but also offer avenues for future targeted therapeutic interventions.

To provide clinical relevance to this issue, we conducted a comprehensive analysis to explore the interplay among gene expression, metabolites, and drug sensitivity across a wide variety of CRC cell lines. Emerging studies have demonstrated that inducing ferroptosis can significantly enhance the effectiveness of cancer cell eradication(Liang et al., 2019). This highlights ferroptosis as an important alternative avenue for cancer treatment. Nonetheless, the precise regulatory mechanisms of ferroptosis in cancer can be complex and require further exploration. We identified numerous anti-cancer drugs that displayed sensitivity to ferroptosis driver genes or resistance in the presence of ferroptosis suppressor genes in a sex-specific manner. For example, we found that *ACSL4*, a ferroptosis driver gene, exhibited an association with the sensitivity of 15 drugs in male KRAS mutant group, including PI3K/mTOR and WNT signaling targeted drugs (**Table S9**).

Currently, there are no large-scale omics studies focused on ferroptosis and CRC that take into account the sex of the patient or KRAS status. Moreover, the importance of ferroptosis induction has not been fully explored in CRC. This has led to a limited understanding of how molecular signatures are affected by mechanisms of ferroptosis and a reasonable interpretation of unexpectedly adverse drug outcomes in anticancer therapy in CRC, particularly in KRAS mutant CRCs. Nonetheless, our study emphasizes the significance of monitoring tumor ferroptosis status in future CRC clinical studies. This extensive study takes into account several factors related to the patient’s cell line origin, including KRAS status, and patient’s sex, making it the most exhaustive study of the ferroptosis landscape in CRC to date. Moreover, we have compiled and classified the drug response status into resistant and sensitive categories, offering a complete overview of the drug response. This detailed mapping of the relationship between molecular features and drug response provides essential biological insights into potential mechanisms of drug insensitivity in some patients, laying a groundwork for future research in this area. The robustness of our findings is confirmed through analysis of multiple large-scale multi-omics datasets, including GSE39582, the TCGA dataset, and an in-house metabolomics cohort.

Our comprehensive analysis emphasizes the potential of precision molecular feature analysis in KRAS mutant CRC tumors. This approach also presents opportunities for drug repurposing (also known as drug repositioning) instead of developing entirely new drugs for specific indications. Considering the significant attrition rates, considerable expenses, and slow process of novel drug discovery and development, the repurposing of ‘old’ drugs for treating both prevalent and rare diseases is gaining traction. By leveraging these findings, we can gain insights into the therapeutic targeting of ferroptosis and explore new possibilities for effective and rapid treatment strategies for sex-specific KRAS-mutant CRC.

## Acknowledgements

This work was supported by funding from the American Cancer Society Research Scholar Grant 134273-RSG-20-065-01-TBE, and the NIH/NCI Yale SPORE in Skin Cancer under award number 5P50CA121974.

## Author contributions

H.Y. conceived the research. C.J. and H.Y. administrated the project and provided guidance on methodology. H.Y. collected GEO dataset and conducted metabolomics data analysis, random forest modeling (VIMP) and drug response analysis. X. S. collected TCGA-COADREAD and conducted survival analysis. S.Y. conducted Gaussian mixture model and Matlab code. S. M. provided guidance on random forest methodology. S. K. provided guidance on CRC clinical information and treatment. H.Y. and X. S. wrote the manuscript and prepared the R code. All authors revised the manuscript.

