## Supplementary figures and tables for "A machine learning and drug repurposing approach to target ferroptosis in colorectal cancer stratified by sex and KRAS": Supplemental Text and Figures.pdf

**Legends of Supplementary Figures and Tables**

**Figure S1. Flowchart of RSF-BE algorithm**. The RSF algorithm adopted a tiered backward elimination approach that adjusts the number of features removed based on the current set's size. If the number of remaining features > M: Remove N at a time. If 50 < remaining features <= M: Remove 10 at a time. If remaining features <= 50: Remove 1 at a time. For mRNA data, M is 200 and N is 100 given a larger feature size. For metabolomics data, M is 100 and N is 30.

**Figure S2. Bootstrapping RSF-BE model performance by sex and KRAS status**. Histogram plots show C-indices with mean, median and standard deviation (SD) from 1000 bootstraps for females and males, females with KRAS MT, females with KRAS WT, males with KRAS MT, and males with KRAS MT from GSE39582, TCGA, and MSKCC cohorts.

**Figure S3. Venn diagrams of RSF-BE results**. A) predictive ferroptosis genes for 5-year OS, B) 5-year PFS, C) predictive metabolites for OS, and D) PFS. Upset plots suggest E) unique predictive genes sets by sex and KRAS for 5-year OS, F) 5-year PFS, and (G) distinct sets of metabolites predictive of OS by females with KRAS MT, females with KRAS WT, males with KRAS MT, and males with KRAS MT.

**Figure S4. Partial dependence plots of predictive genes.** A) males, B) females, C) females with KRAS MT, D) females with KRAS WT, and E) males with KRAS WT from GSE39582.

**Figure S5. Partial dependence plots of predictive metabolites.** A) males, B) males with KRAS MT, C) males with KRAS WT, C) females, B) females with KRAS MT, C) females with KRAS WT in MSKCC cohort.

**Table S1. Demographic tables for GSE39582, TCGA (COADREAD), and MSKCC cohorts.**

**Table S2. Drug information for GDSC**. Sheet I: Drug information of GDSC1; Sheet II: Drug information of GDSC2.

**Table S3. The CCLE CRC cell line information includes KRAS status, sex of patient from whom and tumor type.** Abbreviations: WT: Wild type; MT: Mutant type. In "KRAS" column, "0" presents KRAS wild type, "1" presents exons 1 mutation oncogene KRAS, "2" presents exons 2 mutation oncogene KRAS.

**Table S4. Overlapped genes in male and female CRC patients from TCGA and GSE39582 revealed by variable importance analysis (VMIP).** Sheet I: Overlapped genes in male CRC patients from TCGA and GSE39582 revealed by variable importance analysis (VMIP); Sheet II: Overlapped genes in female CRC patients from TCGA and GSE39582 revealed by variable importance analysis (VMIP).

**Table S5. Overlapped genes in male and female CRC patients revealed by Gaussian mixed model and variable importance analysis (VMIP).** Sheet I: Overlapped genes in male CRC patients revealed by Gaussian mixed model and variable importance analysis (VMIP). Permutation p value and beta value assessing showing variance of genes between KRAS mutant CRCs and KRAS wild-type CRCs. Tumors colon tissues from male patients: KRAS mutant tumors (n = 97), KRAS wild type tumors (n = 166, GSE39582). Sheet II: Overlap genes in female CRC revealed by Gaussian mixed model and variable importance analysis (VMIP). Permutation p value and beta value assessing showing variance of genes between KRAS mutant CRCs and KRAS wild-type CRCs. Tumors colon tissues from female patients: KRAS mutant tumors (n = 86), KRAS wild type tumors (n = 131, GSE39582).

**Table S6. Frequencies of prognostic ferroptosis genes calculated from RSF-BE with 1000 bootstraps for GSE39582 and TCGA, and frequencies of prognostic metabolites calculated from RSF-BE with 1000 bootstraps for MSKCC by sex and KRAS status**. A) OS-related ferroptosis genes, B) PFS-related genes, C) OS- and PFS-related metabolites, D) annotated metabolites list in MSKCC cohort. Green-colored cells in spreadsheet D) indicate overlapping metabolites between OS and PFS.

**Table S7. Aggregate ranks of ferroptosis genes calculated from RSF-BE with 1000 bootstraps by sex and KRAS status.** Green-colored cells indicate overlaps between OS and PFS for each subgroup.

**Table S8. Joint-pathway analysis results by sex and KRAS status on prognostic ferroptosis-related genes and metabolites.** Values in red were significant ($P < 0.05$). Impact scores were colored based on values (darker red means a higher score).

**Table S9. Drug response status across all the GDSC CRC cell lines (Spearman correlation, gene).**

**Table S10. Drug response status across all the GDSC CRC cell lines (Spearman correlation, metabolites).**
