## Supplementary figures and tables for "A machine learning and drug repurposing approach to target ferroptosis in colorectal cancer stratified by sex and KRAS": Supplementary Document.pdf

### Interpretation of High-dimensional Gaussian mixture

Yisha Yao

February 2023

Our goal is to identify a set of genes that are differentially expressed in two groups or a set of genes that would discriminate two groups. Conventionally methods like conducting two-group t test for each individual gene, or Support Vector Machine (SVM) may fail in this specific problem for mainly two reasons. First, there are correlations among the expression levels of these genes, while conducting t tests for individual genes somehow ignores the correlations among the gene expressions. This may lead to spurious detection or fail to detect the genes that truly separate the two groups. Second, SVM involves choosing some kernels that are dependent on some unknown quantities. Different choices of kernels may lead to distinct results.

In this paper, we use a robust and efficient method to identify the set of genes that separate the two groups. We model the expressions of $p$ genes of $n$ individuals by a Gaussian mixture model (Reynolds et al. 2009). Two advantages for using Gaussian mixture model: (1) it is reasonable in biological interpretation; (2) there has been well-established statistical methodology to fit Gaussian mixture model

We first check the multivariate normality of the gene expression data using the R package MVN (Korkmaz et al. 2014). They are approximately distributed as multi-variate Gaussian, and hence it is valid to model the gene expressions by a Gaussian mixture. The rationale behind this idea is as follows. The gene expression patterns of each group follows a multi-variate Gaussian distribution, $N(\boldsymbol{\mu}_1, \boldsymbol{\Sigma})$ and $N(\boldsymbol{\mu}_2, \boldsymbol{\Sigma})$ with the centers $\boldsymbol{\mu}_1$ and $\boldsymbol{\mu}_2$ well-separated. People within the same group have similar but slightly divergent gene expression patterns to account for individual differences.

In the ideal case when the true centers $\boldsymbol{\mu}_1, \boldsymbol{\mu}_2$, covariance $\boldsymbol{\Sigma}$, and component weights $\{w_1, w_2\}$ are known, the optimal classification procedure is the Fisher's linear discriminant rule

$$\mathbf{C}(\boldsymbol{x}) = \begin{cases} 1 & \text{if } (\boldsymbol{x} - \frac{\boldsymbol{\mu}_1+\boldsymbol{\mu}_2}{2})^\top \boldsymbol{\beta} \leq \ln \frac{w_2}{w_1}, \\ 2 & \text{otherwise,} \end{cases} \tag{0.1}$$

where $\boldsymbol{\beta} = \boldsymbol{\Sigma}^{-1}(\boldsymbol{\mu}_1 - \boldsymbol{\mu}_2)$ and $\boldsymbol{x} \in \mathbb{R}^p$ represents the gene expressions of any individual. In our case, $\boldsymbol{\mu}_1, \boldsymbol{\mu}_2, \boldsymbol{\beta}, \{w_1, w_2\}$ are not known and need to be estimated first. After obtaining $\hat{\boldsymbol{\mu}}_1, \hat{\boldsymbol{\mu}}_2, \hat{\boldsymbol{\beta}}, \{\hat{w}_1, \hat{w}_2\}$, we construct the classification rule

$$\mathbf{C}(\boldsymbol{x}) = \begin{cases} 1 & \text{if } (\boldsymbol{x} - \frac{\hat{\boldsymbol{\mu}}_1+\hat{\boldsymbol{\mu}}_2}{2})^\top \hat{\boldsymbol{\beta}} \leq \ln \frac{\hat{w}_2}{\hat{w}_1}, \\ 2 & \text{otherwise,} \end{cases}$$

It has been proved that the above procedure attains minimax optimal error rate in high-dimensional setting $p \gg n$ (Cai et al. 2019). It is easy to obtain $\hat{\boldsymbol{\mu}}_1, \hat{\boldsymbol{\mu}}_2, \{\hat{w}_1, \hat{w}_2\}$, just sample means and frequencies. It takes more efforts to estimate $\boldsymbol{\beta}$. We employ the method proposed in (Cai et al. 2019) to obtain $\hat{\boldsymbol{\beta}}$ because the authors have demonstrated that is the best solution. The detailed procedure is implemented by MATLAB code.

The formula in (0.1) reveals that the quantity deciding the classification of this individual is actually $\boldsymbol{x}\top\boldsymbol{\beta}$, a linear combination of the elements in $\boldsymbol{x}$. Its biological interpretation is that the best "factor" that discriminant two groups is a linear combination of $p$ gene expressions. Indeed, in many cases, there is a set of disease-causing genes. Moreover, some genes might be in the same pathway and interact with each other, so they together lead to some different phenotypes in the two groups.

Since this is a high-dimensional problem (the sample size is smaller than the dimension $p$), the $\hat{\boldsymbol{\beta}} \in \mathbb{R}^p$ we obtained is sparse in the sense that only a small fraction of its elements are nonzero. It means that only the genes corresponding to the nonzero indices of $\hat{\boldsymbol{\beta}}$ are contributing to the classification. In another word, we automatically select a set of discriminating genes through estimating $\boldsymbol{\beta}$.

The levels of metabolites are highly heterogeneous in the sense that some are relatively stable around $10^2$ while some are fluctuating around $10^6$. Such unbalanced variances among different metabolites would result in degenerate multivariate Gaussian distribution. To be more specific: let $\boldsymbol{x} \in \mathbb{R}^p$ be the levels of $p$ metabolites of an individual. $\boldsymbol{x}$ is approximately distributed as a multivariate Gaussian $N(\mathbf{0}, \boldsymbol{\Sigma})$. If $\boldsymbol{\Sigma}$ is positive definite, our method would work well. If $\boldsymbol{\Sigma}$ is not full rank, *i.e.*, the multivariate Gaussian is degenerate, our method would fail. Therefore, we rescale all metabolite levels to remedy the unbalanced variations. It will not change the selection result, *i.e.*, if $x_j$ (the $j$-th metabolite) is contributing to the classification, $\beta_j$ will remain nonzero after rescaling.
