## Supplementary figures and images for "A machine learning and drug repurposing approach to target ferroptosis in colorectal cancer stratified by sex and KRAS"

### Figure S1 COXPH.pdf

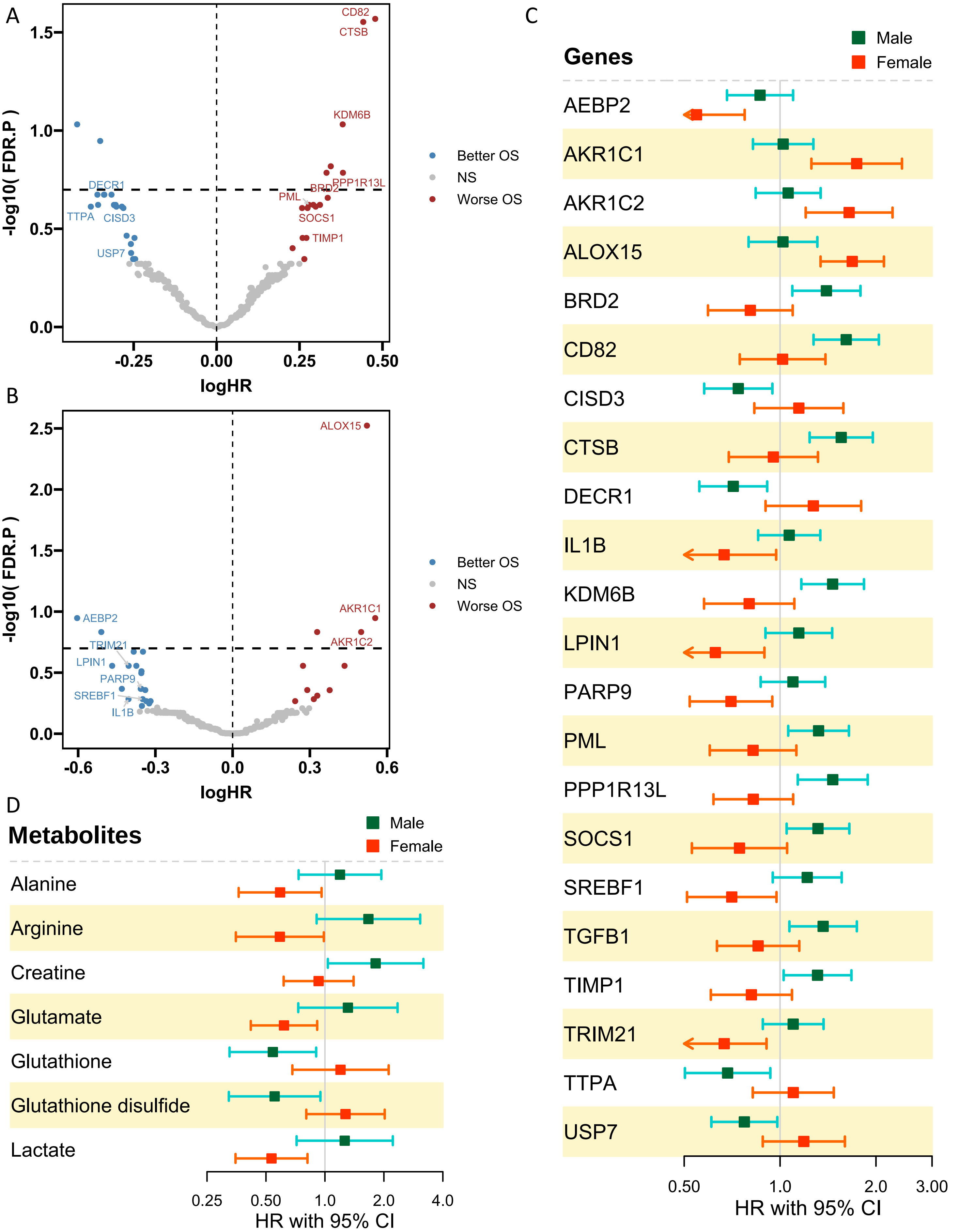

### Figure S1 RSF-BE.pdf

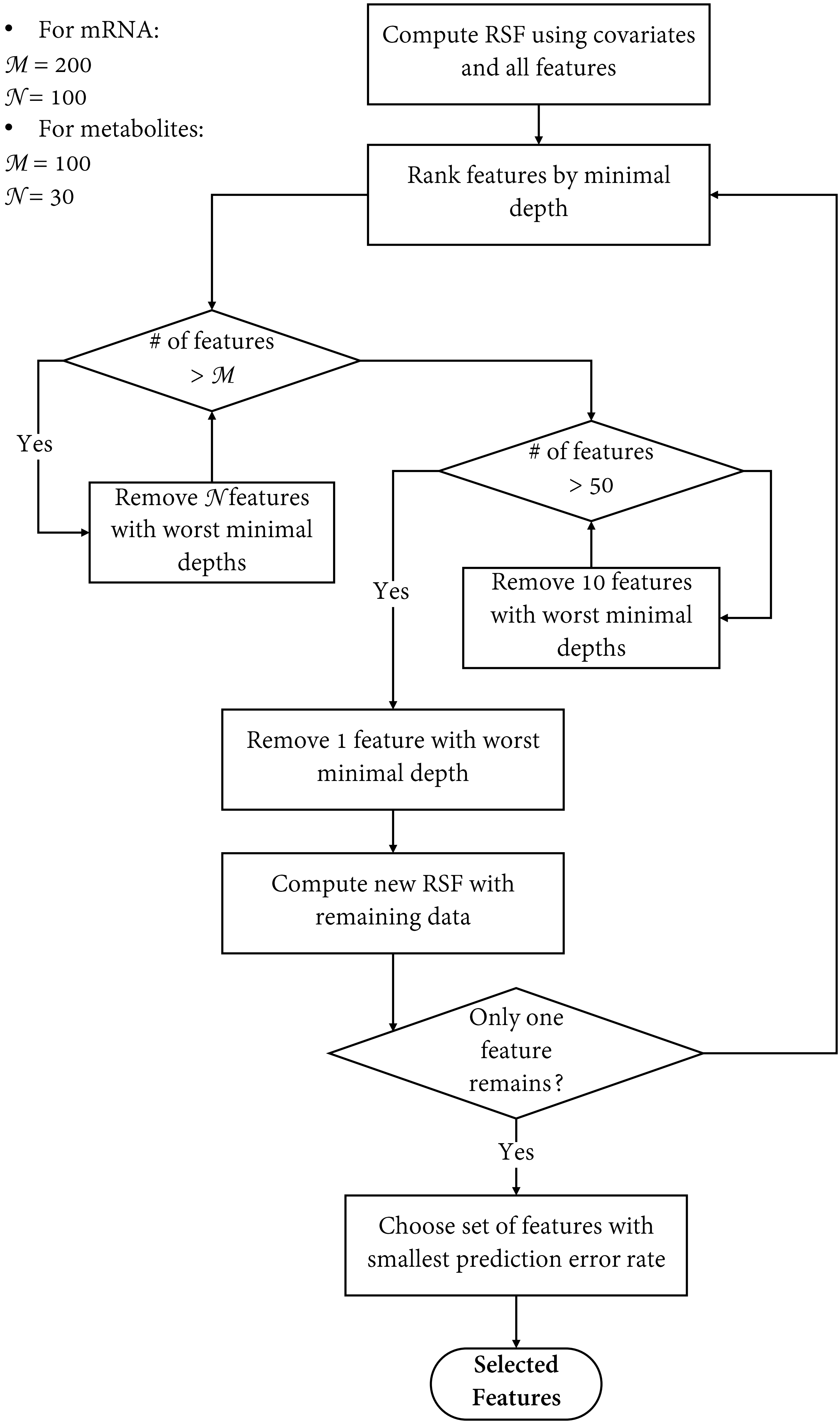

### Figure S2 c_index_T_G_MSKCC_KRAS.pdf

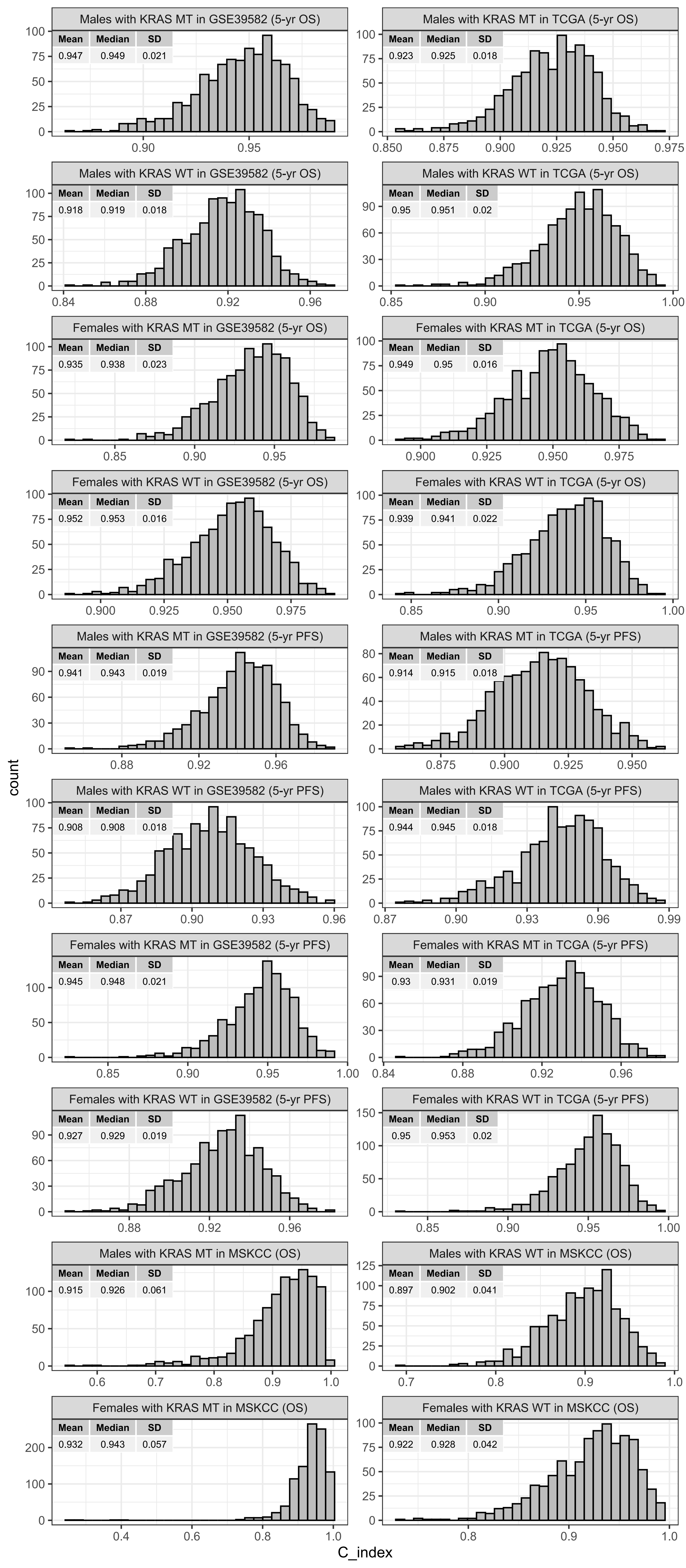

### Figure S3 surv gene mets venn upset.pdf

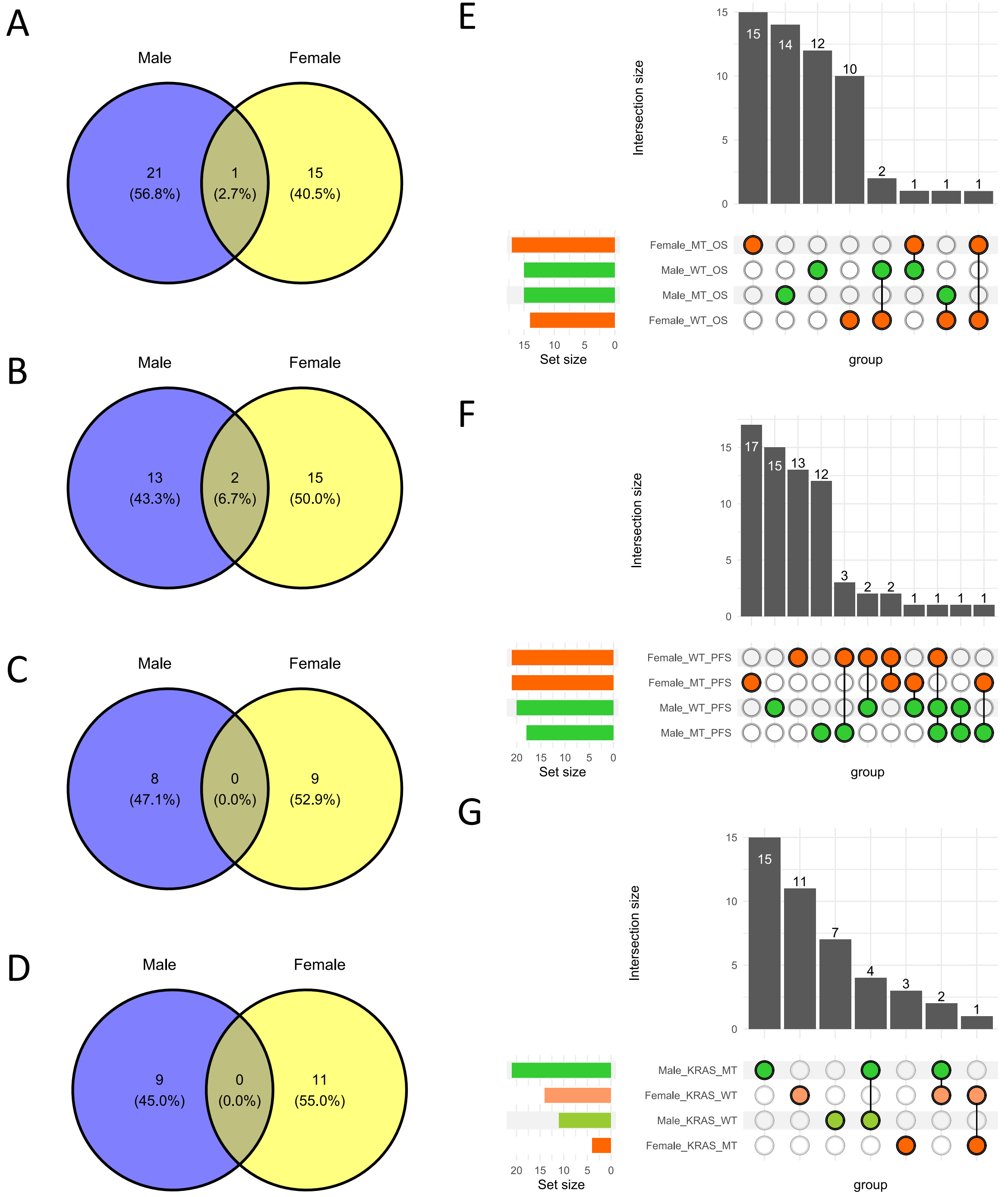

### Figure S4 PartialPlot GEO OS new.pdf

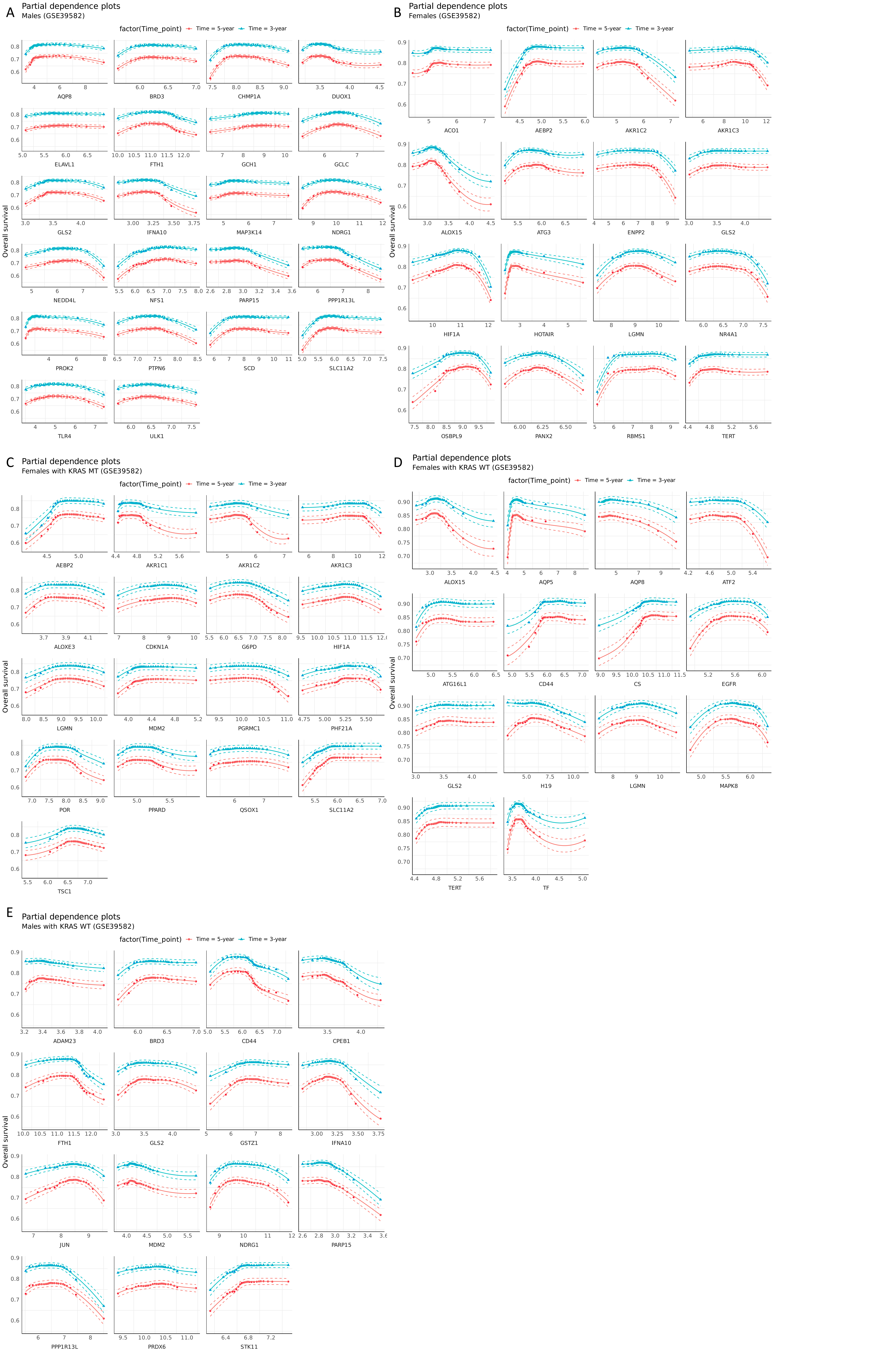

### Figure S5 PartialPlot MSKCC OS.pdf

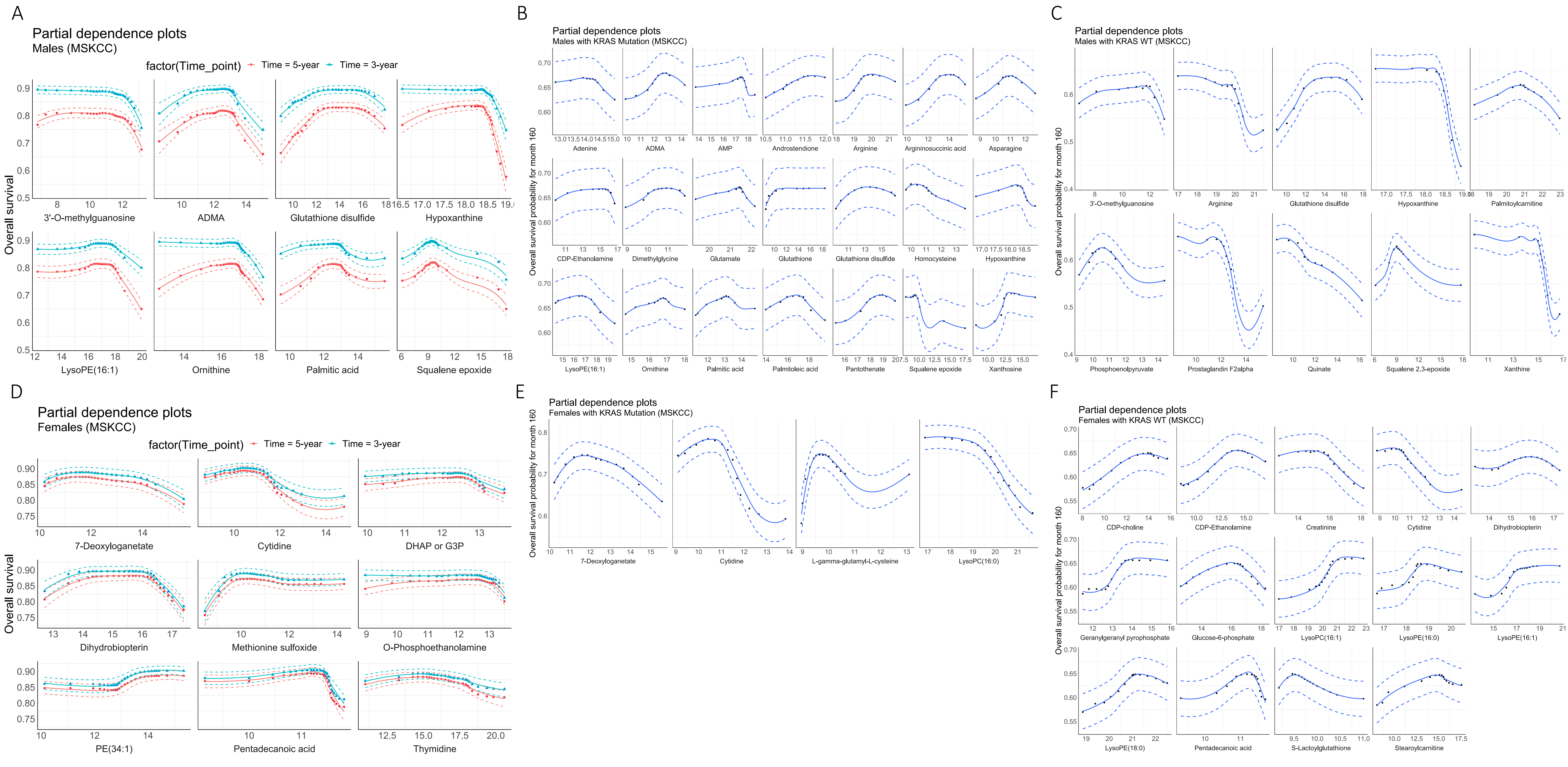
